# Bioinformatic and mechanistic analysis of the palmerolide PKS-NRPS biosynthetic pathway from the microbiome of an Antarctic ascidian

**DOI:** 10.1101/2021.04.05.438531

**Authors:** Nicole E. Avalon, Alison E. Murray, Hajnalka E. Daligault, Chien-Chi Lo, Karen W. Davenport, Armand E.K. Dichosa, Patrick S.G. Chain, Bill J. Baker

## Abstract

Complex interactions exist between microbiomes and their hosts. Increasingly, defensive metabolites that have been attributed to host biosynthetic capability are now being recognized as products of host-associated microbes. These unique metabolites often have bioactivity targets in human disease and can be purposed as pharmaceuticals. Polyketides are a complex family of natural products that often serve as defensive metabolites for competitive or pro-survival purposes for the producing organism, while demonstrating bioactivity in human diseases as cholesterol lowering agents, anti-infectives, and anti-tumor agents. Marine invertebrates and microbes are a rich source of polyketides. Palmerolide A, a polyketide isolated from the Antarctic ascidian *Synoicum adareanum*, is a vacuolar-ATPase inhibitor with potent bioactivity against melanoma cell lines. The biosynthetic gene clusters (BGCs) responsible for production of secondary metabolites are encoded in the genomes of the producers as discrete genomic elements. A candidate palmerolide BGC was identified from a *S. adareanum* microbiome-metagenome based on a high degree of congruence with a chemical structure-based retrobiosynthetic prediction. Protein family homology analysis, conserved domain searches, active site and motif identification were used to identify and propose the function of the ∼75 kb *trans*-acyltransferase (AT) polyketide synthase-non-ribosomal synthase (PKS-NRPS) domains responsible for the stepwise synthesis of palmerolide A. Though PKS systems often act in a predictable co-linear sequence, this BGC includes multiple *trans*-acting enzymatic domains, a non-canonical condensation termination domain, a bacterial luciferase-like monooxygenase (LLM), and is found in multiple copies within the metagenome-assembled genome (MAG). Detailed inspection of the five highly similar pal BGC copies suggests the potential for biosynthesis of other members of the palmerolide chemical family. This is the first delineation of a biosynthetic gene cluster from an Antarctic microbial species, recently proposed as *Candidatus* Synoicohabitans palmerolidicus. These findings have relevance for fundamental knowledge of PKS combinatorial biosynthesis and could enhance drug development efforts of palmerolide A through heterologous gene expression.

## Introduction

Marine invertebrates such as corals, sponges, mollusks, and ascidians, are known to be a rich source of bioactive compounds (Carroll et al., 2019). Due to their sessile or sluggish nature, chemical defenses such as secondary metabolites are often key to their survival. Many compound classes are represented among benthic invertebrates including terpenes, nonribosomal peptide synthetase (NRPS) products, ribosomally synthesized and post-translationally modified peptides (RiPPs), and polyketides. It is estimated that over 11,000 secondary metabolites from marine and terrestrial environments understood to be products of polyketide synthase (PKS) and NRPS origin have been isolated and described (Dejong et al., 2016). BGCs exist as a series of genomic elements that encode for the biosynthetic machinery responsible for production of these secondary metabolites. BGCs can have distinct nucleotide composition properties such as codon usage and guanine-cytosine content that do not match the remainder of the genome (Lawrence et al., 2002; Ravenhall et al., 2015), suggesting a mechanism of horizontal gene transfer from organisms that are distantly related, including across different kingdoms (Schmidt, 2008; Schmitt and Lumbsch, 2009). Interestingly, the BGCs for many natural products isolated from marine invertebrates are found in the host-associated microbiota, reflecting the role of these compounds in symbiosis (Schmidt, 2015).

Polyketides are a complex family of natural products produced by a variety of PKS enzymes that are related to, but evolutionarily divergent from, fatty acid synthases (Helfrich and Piel, 2016). They often possess long carbon chains with varied degrees of oxidation, can contain aromatic components, and may be either cyclic or linear. It is estimated that of the polyketides that have been isolated and characterized, 1% have potential biological activity against human diseases, making this class of compounds particularly appealing from a drug discovery and development standpoint (Koskinen and Karisalmi, 2005). This potential for use as pharmaceuticals is approximately five times greater than for compounds of all other natural product classes (Koskinen and Karisalmi, 2005). Many polyketides are classified as macrolides, which are large-ring lactones that are pharmaceutically relevant due to a number of biological actions, including, targeting the cytoskeleton, ribosomal protein biosynthesis, and vacuolar type V-ATPases (Bordeleau et al., 2005; Nishimura et al., 2005; Napolitano et al., 2012; Ueoka et al., 2015). V-ATPases are responsible for acidification of cells and organelles via proton transport across membranes, including those of lysosomes, vacuoles, and endosomes. These enzymes appear to have an impact on angiogenesis, apoptosis, cell proliferation, and tumor metastasis (Napolitano et al., 2012). A number of marine macrolides inhibit V-ATPases, including lobatamides, chondropsins, iejimalides, and several of the palmerolides (Bowman et al., 2003; Shen et al., 2003; Diyabalanage et al., 2006; Kazami et al., 2006; Noguez et al., 2011).

There are three types of PKS systems. Type I PKS systems in bacteria are primarily comprised of non-iteratively acting multimodular enzymes that lead to progressive elongation of a polyketide chain, though these megaenzymes can also include “stuttering” modules that may act iteratively (Wilkinson et al., 2000; Shen et al., 2007; Tatsuno et al., 2007). In addition, some bacterial Type I PKS systems are comprised solely of iteratively acting monomodular enzymes that catalyze a series of chain elongation steps for polyketide formation (Wang et al., 2020). Type II PKS systems typically contain separate, iteratively acting enzymes that biosynthesize polycyclic aromatic polyketides, while Type III PKS systems possess iteratively-acting homodimeric enzymes that often result in monocyclic or bicyclic aromatic polyketides (Shen et al., 2007). Type I PKS systems can be subdivided into two groups, depending upon whether the acyl transferase (AT) modules are encoded within each module at the site that is parallel to the functional role of the ATs, referred to as *cis*-AT Type I PKS, or physically distinct from the megaenzyme, which are referred to as *trans*-AT Type I PKS. In both cases, there are often parallel relationships between the genome order, the action of enzymatic modules, and the functional groups present in the growing polyketide chain, though in *trans*-AT systems deviations from these parallel relationships is more likely to be observed (Nguyen et al., 2008). In *trans*-AT systems, AT domains may be incorporated in a mosaic fashion through horizontal gene transfer (Nguyen et al., 2008). This introduces greater molecular architectural diversity over evolutionary time, as one clade of *trans*-ATs may select for a malonyl-CoA derivative, while the *trans*-AT domains in another clade may select for unusual or functionalized subunits (Haydock et al., 1995; Jenke-Kodama et al., 2005). Additionally, recombination, gene duplication, and conversion events can lead to further diversification of the resultant biosynthetic machinery (Nivina et al., 2019). Predictions regarding the intrinsic relationship between a secondary metabolite of interest, the biosynthetic megaenzyme, and the biosynthetic gene cluster (BGC) can be harnessed for natural product discovery and development (Kim et al., 2012; Videau et al., 2016; Greunke et al., 2018).

In the search for new and bioactive chemotypes as inspiration for the next generation of drugs, underexplored ecosystems hold promise as biological and chemical hotspots (McClintock et al., 2005). The vast Southern Ocean comprises one-tenth of the total area of Earth’s oceans and is largely unstudied for its chemodiversity. The coastal marine environment of Antarctica experiences seasonal extremes in, for example, ice cover, light field, and food resources. Taken with the barrier to migration imposed by the Antarctic Circumpolar Current and the effects of repeated glaciation events on speciation, a rich and endemic biodiversity has evolved, with consequent potential for new chemodiversity (McClintock et al., 2005; Clarke and Crame, 2010; Young et al., 2013).

Palmerolide A (**Figure 1**) is the principal secondary metabolite isolated from *Synoicum adareanum*, an ascidian which can be found in abundance at depths of 10 to 40 m in the coastal waters near Palmer Station, Antarctica (Diyabalanage et al., 2006). Palmerolide A is a macrolide polyketide that possesses potent bioactivity against malignant melanoma cell lines, while demonstrating minimal cytotoxicity against other cell lines (Diyabalanage et al., 2006). The National Cancer Institute’s COMPARE algorithm was used to correlate experimental findings with a database for prediction of the biochemical mechanism of action by identifying the mechanism of action of palmerolide A as a V-ATPase inhibitor (Paull et al., 1995). Downstream effects of V-ATPase inhibition include an increase in both hypoxia induction factor-1α and autophagy (Diyabalanage et al., 2006; Von Schwarzenberg et al., 2013). Increased expression of V-ATPase on the surface of metastatic melanoma cells (Von Schwarzenberg et al., 2013) perhaps explain palmerolide A’s selectivity for UACC-62 cell lines over all cell types (Diyabalanage et al., 2006). Despite the relatively high concentrations of palmerolide A in the host tissue (0.49–4.06 mg palmerolide A x g^−1^ host dry weight) (Murray et al., 2020), isolation of palmerolide A from its Antarctic source in mass sufficient for drug development it is neither ecologically nor logistically feasible. Although synthetic strategies for palmerolide A have been reported (Jiang et al., 2007; Kaliappan and Gowrisankar, 2007; Nicolaou et al., 2008b; Penner et al., 2009; Lebar and Baker, 2010; Pujari et al., 2011; Pawar and Prasad, 2012; Lisboa et al., 2013), a clear pathway to achieve sufficient quantities needed for drug development has been elusive. Therefore, there is substantial interest in identifying the BGC responsible for palmerolide A production as this would pave a way for future drug development efforts.

**Figure 1.**
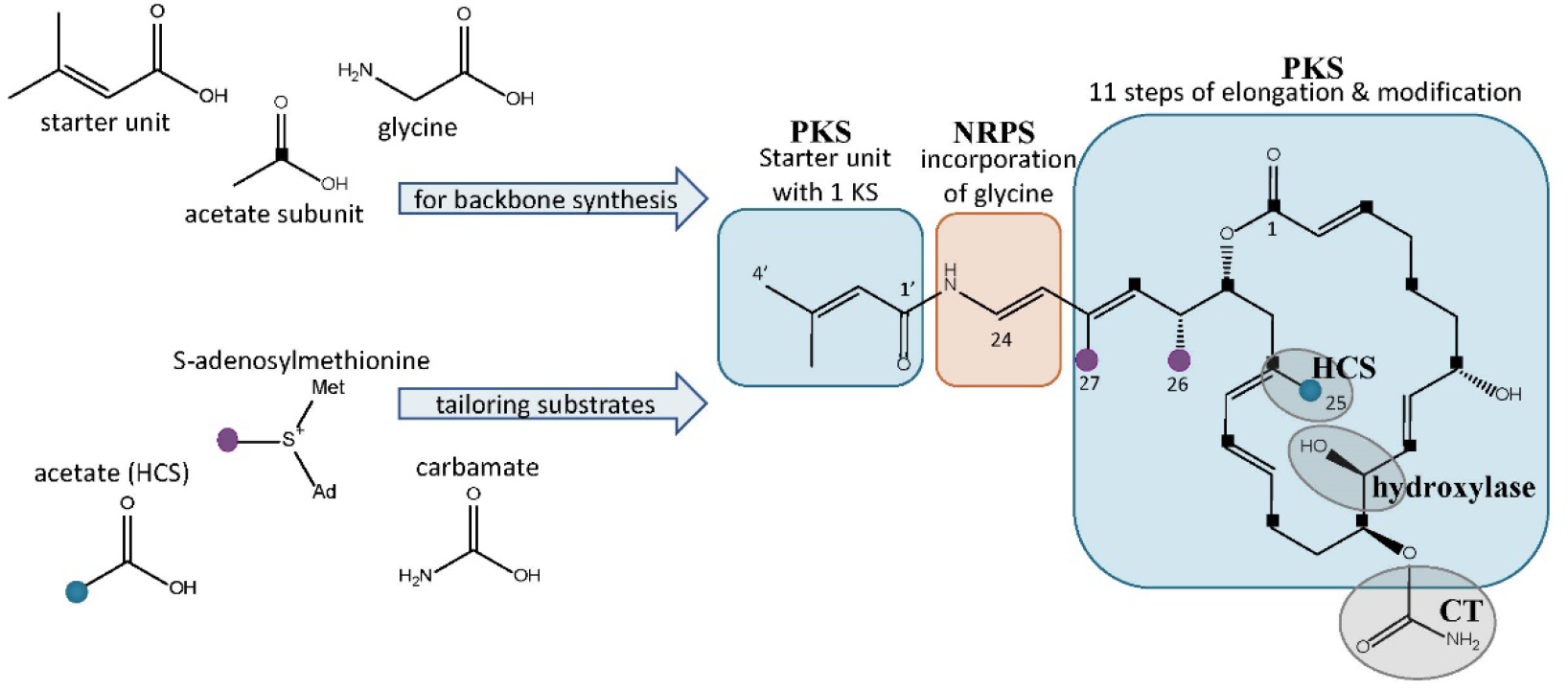
Structure of palmerolide A with notations for the proposed retrobiosynthesis. Backbone synthesis is a result of incorporation of the starter unit, a glycine residue, and acetate subunits (C1 indicated by black squares). Structural features from *trans*-acting tailoring enzymes (indicated by grey ovals) utilize additional substrates: methyl transfers from SAM (purple dots), installation of C-25 methyl from acetate (blue dot) via an HCS cassette, and carbamoyl transfer to the secondary alcohol on C-11. The α-hydroxy group on C-10 is predicted to arise from incorporation of hydroxymalonic acid or a *trans*-acting hydroxylase.

Our approach to identify the palmerolide BGC (*pal* BGC) began with the characterization of the ascidian host-associated microbiome (Riesenfeld et al., 2008). Next, a persistent cohort of bacteria present across many individual ascidians – a core microbiome – for *Synoicum adareanum* was identified through analysis of occurrence of distinct amplicon sequence variants (ASV) from iTag sequencing of the Variable 3-4 regions of the bacterial 16S rRNA (Murray et al., 2020). This work ultimately led to the evaluation of the microbiome metagenome and the subsequent assembly of a metagenome assembled genome (MAG) of *Candidatus* Synoicohabitans palmerolidicus, a verrucomicrobium in the family *Opitutaceae* (Murray et al., 2021). Contained within the genome are five non-identical copies of a candidate *pal* BGC. Here, we report on a detailed bioinformatic analysis of the *pal* BGCs and conclude that at least three of the candidate BGCs likely are responsible for the biosynthesis of palmerolides with structures that have been previously reported from Antarctic *S. adareanum* in this macrolide family (Diyabalanage et al., 2006; Noguez et al., 2011).

## Materials and Methods

The details of field collections, sample processing and genomic methods (sequencing, assembly, and analysis) used to identify the putative biosynthetic gene clusters are described in detail in two related publications (Murray et al., 2020, 2021). In brief, *Synoicum adareanum* samples were collected by SCUBA from the Antarctic Peninsula in the Anvers Island Archipelago and flash frozen. Microbial cells were separated from host tissue using a homogenization protocol established for this holobiont followed by differential centrifugation to separate the host cellular debris and microbial cells (Murray et al., 2020). The metagenome assembled genome (MAG) sequence analyzed in this study was generated from two *S. adareanum* samples (Bon-1c-2011 and Del-2b-2011) with high copy numbers of the putative biosynthetic gene cluster sequenced using PacBio technology. The resulting assembly of produced a nearly complete 4.3 MB genome, with five unique contigs and five varying copies of the *pal* BGC (referred to as *pal* BGC 1 through *pal* BGC 5) (Murray et al., 2021). Given the phylogenomic novelty of the MAG, the name *Candidatus* Synoicihabitans palmerolidicus was proposed for this new genus in the *Opitutaceae* family (Verrucomicrobia phylum).

The methods employed in this study used bioinformatic tools to develop predictive models of palmerolide biosynthesis. Enzymatic reactions and organic synthetic interpretations were based on homology analyses. Automated annotation and manual bioinformatic tools were used to discern the details of palmerolide A biosynthesis in addition to generating predictions for the other BGCs. The *Ca*. S. palmerolidicus MAG was annotated using antiSMASH (v. 5.0) (Blin et al., 2019) using the full complement of annotation options available. Then we predicted the gene cluster responsible for palmerolide A biosynthesis using retrobiosynthetic predictions focused on the 5’ end of the BGCs (**Figure 1**). Only one of the five BGCs met the criteria for a non-ribosomal peptide glycine starter unit. The annotation predictions were integrated and validated with results of additional protein family homology analysis, conserved domain searches, active site and motif identification to predict the step-wise biosynthesis of palmerolide A. Manual annotation of the *pal* BGC sequences included BLASTP searches to confirm enzymatic identities, then protein family alignments were used to identify active site residues key for stereochemical outcomes, confirm substrate affinities, and other biochemical synthesis details.

Additional manual bioinformatic efforts included obtaining BGCs from public NCBI databases for basiliskamide, bryostatin 1, calyculin, corallopyronin, mandelalide, onnamide, oxazolamycin, pederin, phormidolide, psymberin, sorangicin, and myxoviricin (**Supplemental Table S2**). ClustalO alignment tool in the CLC Genome Workbench (QIAGEN aarhau A/S v. 20.0.3) was used for multiple sequence alignments of enzymatic domains with HMM Pfam Seeds obtained from EMBL-EBI and the amino acid sequences from the other PKS BGCs. MIBiG (Kautsar et al., 2020) was used to acquire the KS amino acid sequence from the type III PKS BGC responsible for 3-(2’- hydroxy-3’-oxo-4’-methylpentyl)-indole biosynthesis from *Xenorhabdus bovienii SS-2004* (GenBank Accession: FN667741.1), which was used for an outgroup. The *pal* BGC ACPs and PCPs were numbered according to their position in the proposed biosynthesis of palmerolide A (**Supplemental Figures S1 and S2**). The BGC KSs were numbered according to their position in their proposed biosynthesis in the literature. Prior to the construction of the phylogenetic tree for the KS domains (**Supplemental Figure S4**), the sequences in the alignment were manually inspected and trimmed. Phylogenetic trees were created in the CLC Genome Workbench (QIAGEN aarhau A/S v. 20.0.3) with Neighbour Joining (NJ) as a distance method and Bayesian estimation for ACP and PCP comparisons as well as for KS analysis. Jukes-Cantor was selected for the genetic distance model and bootstrapping with performed with 100 replicates. Additionally, the sequence of each KS in the *pal* BGCs was queried using the *trans*-AT PKS Polyketide Predictor (*trans*ATor) to help define the specificity of KS domains. The software is based on phylogenetic analyses of fifty-four *trans*-AT type I PKS systems with 655 KS sequences and the resulting clades are referenced to help predict the KS specificity for the upstream unit (Helfrich et al., 2019).

## Results and Discussion

### A. Retrobiosynthetic Scheme for Palmerolide A

A retrobiosynthetic scheme of the *pal* BGC was developed based on the chemical structure for palmerolide A, including modules consistent with a hybrid PKS-NRPS with tailoring enzymes for key functional groups (**Figure 1**). We hypothesized that the initial module would be PKS-like in nature to utilize 3-methylcrotonic acid as the starter unit followed by a NRPS domain for the incorporation of glycine. PKS elongation was predicted to be an 11-step sequence resulting in 22 contiguous carbons. Modifying enzymes that are encoded co-linearly were predicted to create the architectural diversity with olefin placement, reduction of certain carbonyl groups to secondary alcohols, and full reduction of other subunits. In addition, incorporation of methylmalonyl CoA or enzymatic activity of carbon methyltransferases (cMTs) were predicted to be responsible for the placement of methyl groups C-26 and C-27 from *S*-adenosylmethionine (SAM).

Several key structural features proposed to result from the action of *trans*-acting enzymes are present. For example, as seen in the kalimantacins (Mattheus et al., 2010), the carbamate on C-11 was hypothesized to be installed by a carbamoyl transferase (CT). The C-25 methyl group located on C-17 in the β-position to the carbonyl suggests the origin of this branch is likely from hydroxymethylglutaryl-CoA synthase (HCS) catalysis, rather than SAM-mediated methylation, which occurs at the α-position to the carbonyl. SAM-mediated methylation does however appear to be the origin of the C-26 and C-27 methyl groups. Methylation in this acetate carboxyl position is unusual, but represented in a number of notable BGC’s, such as those of the jamaicamides, bryostatins, curacin A, oocydin, pederin, and psymberin, among others; in biochemically characterized Type I PKS BGCs, HCS-mediated β-branch formation is the common mechanism (Chang et al., 2004; Edwards et al., 2004; Sudek et al., 2007; Fisch et al., 2009; Matilla et al., 2012). Lastly, the hydroxy group on C-10 in the α-position of the acetate subunit, was hypothesized to arise by elongation resulting from hydroxymalonyl-CoA incorporation or by the action of a hydroxylase at a later stage of biosynthesis.

### B. Proposed Architecture of the Putative *pal* Biosynthetic Gene Cluster and Biosynthesis of Palmerolide A

The *Ca*. Synoicohabitans palmerolidicus MAG reported a candidate hybrid PKS-NRPS biosynthetic gene cluster that was present in multiple, non-identical copies (Murray et al., 2021). Detailed inspection of one of these clusters (specifically contig 9 which corresponds to *pal* BGC4, the first to be interrogated here) has excellent congruence with the retrobiosynthetic predictions outlined above (**Figure 1**). The results here in which we integrated BGC annotations predicted using AntiSMASH (Blin et al., 2019) with information from protein family homology analysis, conserved domain searches, active site and motif identification, together support the hypothesis that this 75 kb BGC is putatively responsible for palmerolide A production.

The architecture of the biosynthetic gene cluster reveals core biosynthetic domains followed by 2 ATs, and finally, a series of trans-acting domains (**Figure 2**). The core scaffold is explained by the NRPS and *trans*-AT PKS hybrid system. In addition, each of the tailoring enzymes that are expected for the distinct chemical features are encoded in the *Ca*. Synoicihabitans palmerolidicus genome. Comparisons of this overall modular architecture with eleven other *trans*-AT systems suggests a significant amount of flexibility in these constructs (**Figure 2**). The mandelalide BGC (Lopera et al. 2017) most closely resembles that of palmerolide in which the core modules are followed in line by AT modules, and trans-acting modules are encoded at the end of the cluster, except that there is only a single AT reported in the case of mandelalide. The proposed BGC for palmerolide A is comprised of 14 core biosynthetic modules and 25 genes in a single operon of 74,655 bases (**Figure 3**). Fourteen modules are co-linear and two *trans*-AT domains (modules 15 & 16) follow the core biosynthetic genes. Additional *trans*-acting genes contribute to backbone modifications with at least one gene contributing to post-translational tailoring (**Figure 3**).

**Figure 2.**
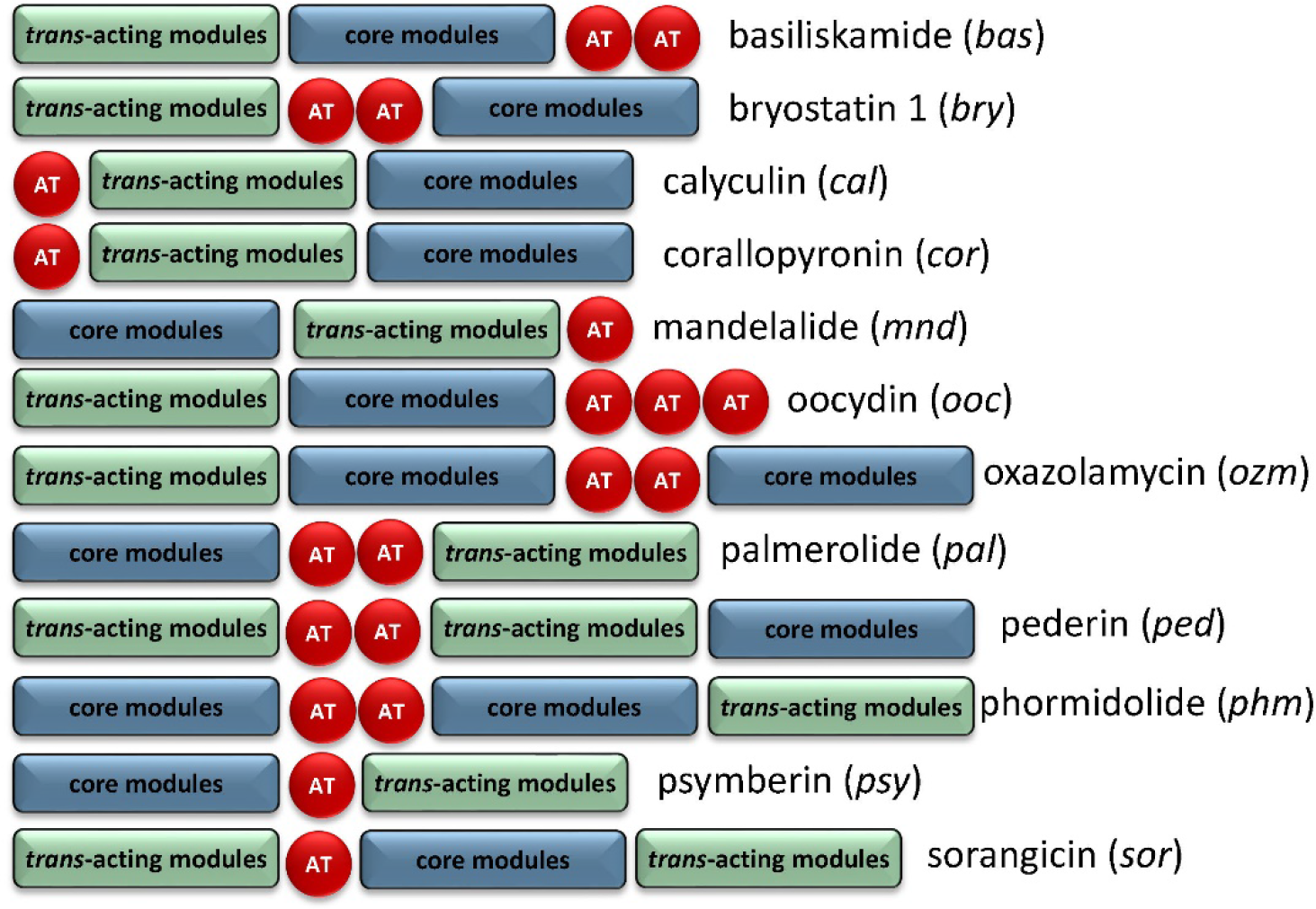
Comparison of BGC organization of select *trans*-AT systems. There is significant variability in the order of the core modules, AT modules, and modules which contain *trans*-acting tailoring enzymes. There is also variability in the number of encoded AT modules, though the AT modules are typically encoded on separate, but tandem genes if more than one is present.

**Figure 3.**
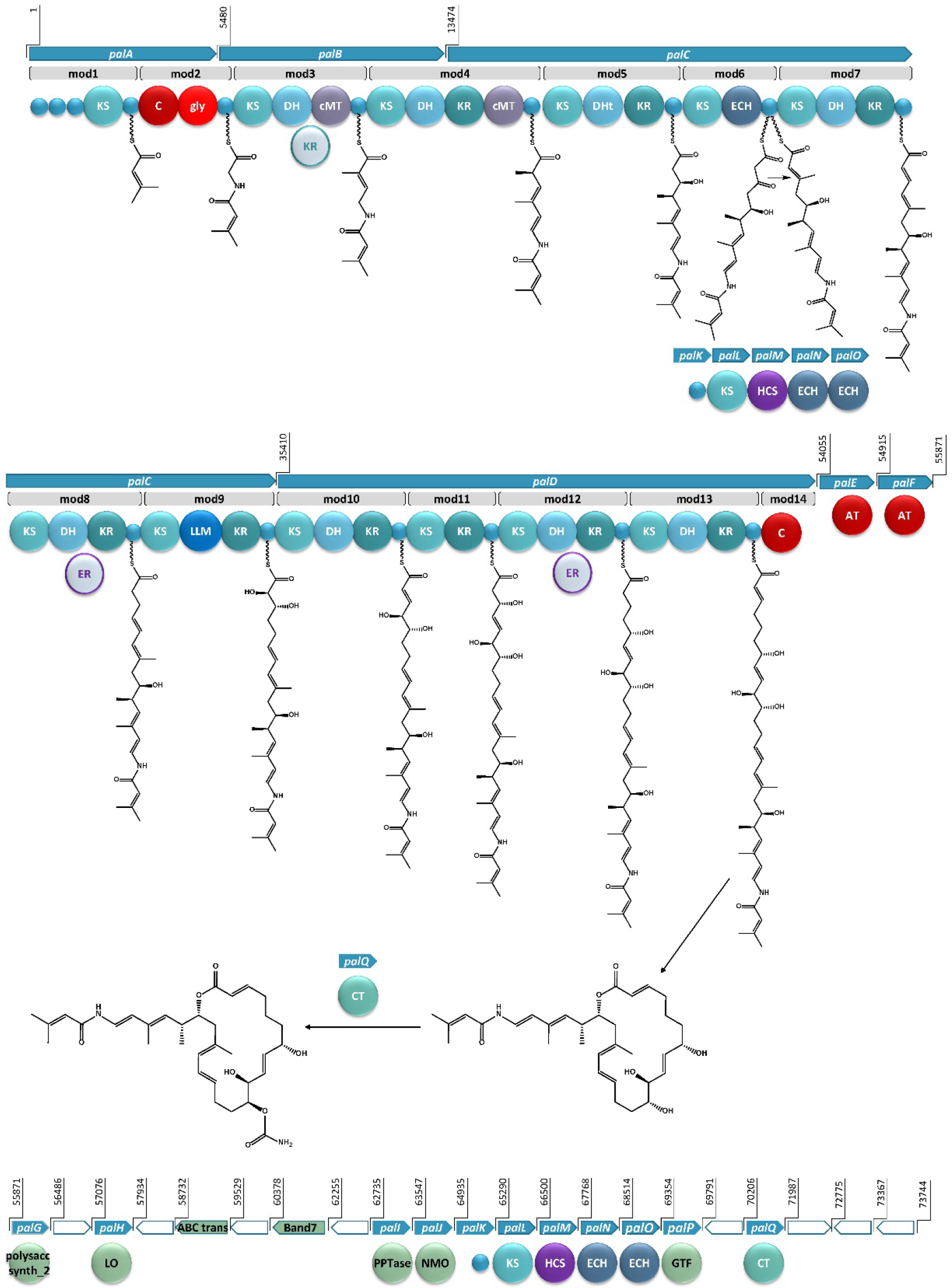
The proposed BGC for palmerolide A, showing the hybrid PKS-NRPS system. KS: ketosynthase domain, C: condensation domain, gly: adenylation domain for glycine incorporation, DH: dehydratase domain, cMT: carbon methyl transferase domain, KR: ketoreductase domain, DHt: dehydratase variant; ECH: enoyl-CoA hydratase, LLM: luciferase-like monooxygenase, AT: acyl transferase; polysacc synt_2: polysaccharide biosynthesis protein, LO: lactone oxidase, ABC trans: ATP-binding cassette transporter, Band7: stomatin-like integral membrane, PPTase: phosphopantetheinyl transferase, NMO: nitronate monooxygenase, HCS: hydroxymethylglutaryl-CoA synthase, GTF: glycosyl transferase ER: enoyl reductase, CT: carbamoyl transferase, small blue circles represent acyl- or peptidyl-carrier proteins. Ppant arms are symbolized by wavy lines. The grey domains indicate domains that would be expected to perform an enzymatic transformation; however, are not encoded in the BGC. Blue arrows indicate biosynthetic genes. Green arrows indicate genes that encode for non-biosynthetic proteins. White arrows reflect hypothetical genes. The BGC is displayed in reverse compliment.

#### An unusual starter unit and NRPS domains of *palA*

Bioinformatic analysis of the gene sequence suggests that the initial core biosynthetic domains of *palA* (modules 1 and 2) encode for the requisite acyl carrier proteins (ACP) (**Figure 3**). ACPs are typically responsible for tethering the acyl subunits to a phosopantetheine arm via thioester bond formation. Encoded in module 1 are three ACPs in tandem, which could serve to promote an increase in metabolite production (Gulder et al., 2011). The second in series is an ACP-β containing the conserved domain sequence GXDS (Bertin et al., 2016) which is likely the acceptor of a starter unit containing a β-branch. This is consistent with our proposed starter unit for palmerolide A, 3-methylcrotonic acid. While both *trans*-acting ATs, PalE and PalF, (**Figure 3**) possess the catalytic active site serine which is key for the proper positioning of the selected subunit within the hydrophobic cleft of the active site (Reeves et al., 2001; Helfrich and Piel, 2016), only the first AT, PalE, has a characteristic motif that includes an active site phenylalanine, conferring specificity for malonate selection (Yadav et al., 2003). The AT selecting the methylcrotonic acid starter unit is likely the second of the two *trans*-AT domains (PalF), which lacks definitive specificity for malonyl-CoA. In support of this hypothesis, some *trans*-acting ATs have demonstrated affinity for a wider range of substrates than their *cis*-acting counterparts (Dunn et al., 2014; Nivina et al., 2019). 3-methylcrotonyl-CoA is an intermediate of branched-chain amino acid catabolism in leucine degradation; intermediates of this pathway can be diverted to secondary metabolite production (Díaz-Pérez et al., 2016). Interestingly, an upregulation of leucine catabolic pathways can be seen as a mechanism of cold adaption as intermediates, such as methylcrotonyl-CoA can be incorporated into the biosynthetic pathways of branched chain fatty acids (Mahmud et al., 2002; Díaz-Pérez et al., 2016). Integration of branched chain fatty acids in cell walls allows some microorganisms to maintain cell wall fluidity during temperature fluctuations (Klein et al., 1999; Fonseca et al., 2011, 2013), a factor which may be at play in this Antarctic bacterium. The subsequent NRPS module (module 2) contains condensation (C) and adenylation (A) domains as well as a carrier protein. Signature sequence information and NRPSPredictor2 analysis (Röttig et al., 2011; Blin et al., 2019) of the A domain are consistent with selection of a glycine residue. These domains incorporate the amino acid residue, resulting in the addition of a nitrogen and two carbons in this step of the biosynthesis of palmerolide A.

In a non-canonical fashion, the carrier proteins flanking the NRPS domains do not appear to be the expected ACP and peptidyl-carrier protein (PCP) for module 1 and 2, respectively. The carrier protein following the KS domain in module 1 was initially annotated as a non-β-branching ACP, however, phylogenetic analysis with the amino acid sequences of carrier proteins from other hybrid PKS-NRPS systems demonstrates that this carrier protein is in the same clade as PCPs (**Supplemental Figure S1**). The carrier protein associated with module 2, which was initially annotated simply as a phosphopantetheine attachment site (Pfam00550.24), is found to be more phylogenetically-related to ACPs within PKS-NRPS systems (**Supplemental Figure S1**). Notably, it possesses the (D/E)xGxDSL motif for phosphopantetheine arm attachment (Keatinge-Clay, 2012) with the exception of an isoleucine rather than leucine in the final position of the motif, which is a residue common to other ACPs from hybrid PKS-NRPS systems (**Supplemental Figure S2**). Typically, a PCP would follow the domains in NRPS-like modules, however, there are exceptions in the literature. For example, the BGCs for both corallopyronin and oxazolamycin contain ACPs following an A domain (Erol et al., 2010; Zhao et al., 2010). This non-canonical finding could point to the acquisition of these domains over evolutionary time, as the carrier protein for module 1 is encoded in *palA*, the same gene encoding the proteins for both modules 1 and 2, whereas the carrier protein for module 2 is encoded at the beginning of *palB*, a gene which encodes for only PKS domains (**Figure 3**).

#### Contiguous PKS chain and *trans*-acting enzymes at site of action for *palB* – *palD*

The contiguous carbon backbone of palmerolide A arises from 11 cycles of elongation in which the synthesis proceeds through a series of modules with a variety of enzymatic domains that include an ACP, KS, and associated genes that establish the oxidation state of each subunit (**Figure 3**). The first module of *palB* (module 3) includes a dehydratase (DH) and cMT domains, a sequence which results in a chain extension modification to an α,β-unsaturated thioester, a result of the action of the encoded DH. The expected KR domain that would be responsible for the Δ^22^ olefin (**Figure 3**) is not present. The BGCs for bryostatin 1, corallopyronin, and sorangicin also lack an accompanying KR domain to work in concert with an encoded DH. The unaccompanied DH in the bryostatin 1 and corallopyronin systems are deemed inactive; however, an olefin results from the DH in the absence of an accompanying KR in both modules 9 and 11 of the sorangicin BGC (Sudek et al., 2007; Erol et al., 2010; Irschik et al., 2010). The subsequent cMT methylation is consistent with an *S*-adenosylmethionine (SAM)-derived methyl group, as expected for C-27 in palmerolide A. Module 4, spanning the end of *palB* and beginning of *palC*, includes a DH, a ketoreductase (KR), and another cMT domain. This cluster of domains is predicted to result in the methyl-substituted conjugated diene of the macrolide tail (C-19 through C-24, C-26 on palmerolide A).

The substrate critical for macrolactonization of the polyketide is the C-19 hydroxy group, a result of the action of the KS and KR domains encoded in module 5 (**Figure 3**). Interestingly, a domain initially annotated as a dehydratase (DHt) at this location may contribute to the final cyclization and release of the molecule from the megaenzyme by assisting the terminal C domain with ring closure (Bertin et al., 2016). The DHt sequence does not possess the hotdog fold that is indicative of canonical dehydratases (Cantu et al., 2010), and therefore, may not truly represent a DH. Alternatively, this domain could be responsible for the olefin shifts to the β,γ-positions, as seen in bacillaene and ambruticin biosynthesis (Moldenhauer et al., 2010; Berkhan et al., 2016).

In addition to a standard ACP and KS encoded in module 6, which would lead to a ketone function, an enoyl-CoA hydratase (ECH) is also encoded. Based on our retrobiosynthetic analysis, the ketone at C-17 is the necessary substrate for HCS-catalyzed β-branch formation, resulting in the C-25 methyl group on C-17. We propose that the ECH encoded in module 6 works in concert with the HCS cassette. The HCS cassette (PalK through PalO) is comprised of a series of *trans*-acting domains, including an ACP, an HCS, a free KS, and 2 additional ECH modules (**Figure 3**). The HCS cassette can act while the elongating chain is tethered to an ACP module, rather than after cyclization and release (Moldenhauer et al., 2007; Hertweck, 2009). The two ECHs in the HCS cassette along with the ECH encoded in-line with the core biosynthetic genes would be responsible for isomerization of a terminal methylene to the observed internal olefin. An HCS cassette formed by the combination of a *trans*-KS and at least one ECH module with an HCS domain is reported in several other bacterial BGCs such as bryostatin 1, calyculin A, jamaicamide, mandelalide, phormidolide, and psymberin (Edwards et al., 2004; Esquenazi et al., 2008; Fisch et al., 2009; Wakimoto et al., 2014; Bertin et al., 2016; Lopera et al., 2017). The domain structure for the HCS cassettes has a remarkably high degree of synteny across these diverse BGCs (Buchholz et al., 2010), however, the presence of an in-line ECH domain in these biosynthetic systems may vary. There is precedence for a similar domain architecture in oocydin, pederin, onnamide, psymberin, phormidolide, and mandelalide, though the presence of the additional ECH domain in-line with the core biosynthetic genes does not necessarily correlate with the formation of an internal versus terminal olefin (Piel et al., 2004; Fisch et al., 2009; Matilla et al., 2012; Bertin et al., 2016; Lopera et al., 2017).

There is substantial similarity in the domain structure of module 7, module 10, and module 13, whereby each includes a KS, DH, and KR (**Figure 3**). The olefin that arises from the action of module 7, concomitant with carbon chain elongation, is conjugated with the Δ^16^ olefin adjacent the C-17 β-branch. Modules 10 and 13 have similar enzymatic composition to 7 and are likely responsible for Δ^8^ and Δ^2^ olefins. The combination of KR and DH domains are also found in modules 8 and 12; however, in concert with an as of yet unidentified *trans*-acting enoyl reductase domain (ER), these olefins would be reduced to fully saturated monomeric subunits. There are some examples of *trans*-acting ER domains carrying out this function, including OocU in oocydin, SorN in sorangicin, and MndM in mandelalide (Irschik et al., 2010; Matilla et al., 2012; Lopera et al., 2017), while in other systems, such as corallopyronin and leinamycin, the reductions of the olefins are largely unexplained (Cheng et al., 2003; Erol et al., 2010). The reduction by a *trans*-acting enzyme often occurs while the elongating polyketide is tethered to the megaenzyme, as evidenced by the downstream specificity of the KS module for Claisen-type condensation with subunits containing single or double bonds (Irschik et al., 2010).

The genetic architecture for the biosynthesis of two functional groups essential for bioactivity is encoded in module 9 (**Figure 3**). Structure-activity relationship studies demonstrate the importance of the C-10 hydroxyl group and the C-11 carbamate (Nicolaou et al., 2008a). The KR domain predicting the C-11 alcohol function serves as the substrate for the carbamoyl transferase (*palQ*) in a post-translational modification (Haydock et al., 2005; Chen et al., 2009; Mihali et al., 2011). Intriguingly, a domain annotated as a luciferase-like monooxygenase (LLM) initially seems out of place. However, palmerolide A has a hydroxy group at C-10, which is the α-position of the acetate subunit inserted in module 9. LLMs associated with BGCs may not serve as true luciferases, but, instead, demonstrate oxidizing effects on polyketides and peptides without evidence of corresponding bioluminescence (El-Sayed et al., 2001; Maier et al., 2015). For example, there is an overrepresentation of LLMs in *Candidatus* Entotheonella BGCs without known bioluminescence (Lackner et al., 2017). As demonstrated through individual inactivation of the LLM in the BGC of mensacarin, a Type II PKS system, Msn02, Msn04, and Msn08 have key activity as epoxidases and hydroxlases (Maier et al., 2015). There are several examples of LLMs in modular Type I PKS systems. OnnC from onnamide and NazB from nazumamide are two LLMs in *Candidatus* Endotheonella that are proposed to serve biosynthetically as hydroxylases (Lackner et al., 2017). In calyculin and mandelalide, the CalD and MndB LLMs catalyze chain shortening reactions through α-hydroxylation and Baeyer-Villiger-type oxidation reactions (Wakimoto et al., 2014; Lopera et al., 2017). Phormidolide has a LLM that adds a hydroxy group, which is hypothesized to attack an olefin through a Michael-type addition for cyclization with enzymatic assistance from a pyran synthase (Bertin et al., 2016). The hydroxylation that is key in cyclization of oocydin A is likely installed by OocK or OocM, flavin-dependent monooxygenases that are contiguous to the PKS genes and are thought to act while the substrate is bound to a portion of the PKS megaenzyme (Matilla et al., 2012). It is this hydroxylase activity that we propose for the LLM in module 9. Since the producing bacteria is yet to be cultured, it is not established whether this LLM may also serve a role in bioluminescence and/or quorum sensing. Further evidence for the role of the LLM is provided through alignment against other LLM’s. The *pal* BGC contains both LuxR family transcriptional regulator as well as the DNA-binding response regulator. In addition to the annotation within Pfam00296, which includes the bacterial LLMs, the sequence, of which, is homologous with the multiple sequence alignments and the hidden Markov models of the TIGR subfamily 04020 of the conserved protein domain family cl19096 noted for natural product biosynthesis LLMs (Lackner et al., 2017). The subfamily occurs in both NRPS and PKS systems as well as small proteins with binding of either flavin mononucleotide or coenzyme F420. Alignment of the LLMs from multiple PKS systems, including palmerolide A, shows homology with model sequences from the TIGR subfamily 04020 (**Supplemental Figure S3**).

The addition of C-5 and C-6 and the reduction of the β-carbonyl to form the C-7 hydroxy group of palmerolide A, is due to module 11, which possesses a KR domain in addition to elongating KS (**Figure 3**). In the structure of palmerolide A this is followed by the fully reduced subunit from module 12 as discussed above. The final elongation results from module 13, which includes DH and KR domains that contribute to the conjugated ester found as palmerolide A’s C-1 through C-3, completing the palmerolide A C_24_ carbon skeleton.

#### Noncanonical termination condensation domain in *palD* for product cyclization and release

Typically, PKS systems terminate with a thioesterase (TE) domain, leading to release of the polyketide from the megaenzyme (Piel, 2002; Gu et al., 2009; Gehret et al., 2011; Lopera et al., 2017). This canonical domain is not present in the *pal* cluster. Instead, the final module in the *cis*-acting biosynthetic gene cluster includes a truncated condensation domain comprised of 133 amino acid residues, compared to the approximately 450 residues that comprise a standard condensation domain (Stachelhaus et al., 1998) (**Figure 3**). Condensation domains catalyze cyclization through ester formation in free-standing condensation domains that act in *trans* as well as in NRPS systems (Zaleta-Rivera et al., 2006; Lin et al., 2009). In addition, this non-canonical termination domain is not without precedent in hybrid PKS-NRPS and in PKS systems as both basiliskamide and phormidolide include condensation domains for product release (Theodore et al., 2014; Bertin et al., 2016). Though the terminal condensation domain in the *pal* BGC is shortened, it maintains much of the HHXXDDG motif (**Supplemental Figure S3**), most notably the second histidine, which serves as the catalytic histidine in the condensation reaction (Stachelhaus et al., 1998).

#### Stereochemical and structural confirmation based on sequence information

KR domains are NADPH-dependent enzymes that belong to the short-chain dehydrogenase superfamily, with Rossman-like folds for co-factor binding (Keatinge-Clay and Stroud, 2006; Keatinge-Clay, 2012). Enzymatically, the two KR subtypes, A-Type and B-type, are responsible for stereoselective reduction of β-keto groups and can also determine the stereochemistry of α-substituents. C-type KRs, however, lack reductase activity and often serve as epimerases. A-Type KRs have a key tryptophan residue in the active site, do not possess the LDD amino acid motif, and result in the reduction of β-carbonyls to L-configured hydroxy groups (Keatinge-Clay, 2012). B-Type, which are identified by the presence of an LDD amino acid motif, result in formation of D-configured hydroxy groups (Keatinge-Clay, 2012). The stereochemistry observed in palmerolide A is reflected in the active site sequence information for the D-configured hydroxyl groups from module 5 and module 11 (**Figures 1** and **3**). When an enzymatically active DH domain is within the same module, the stereochemistry of the *cis*-versus *trans*-olefin can be predicted, as the combination of an A-Type KR with a DH results in a *cis*-olefin formation and the combination of a B-Type KR with a DH results in *trans*-olefin formation. The *trans*-α,β-olefins arising from module 7 (Δ^14^), module 10 (Δ^8^), and module 13 (Δ^2^) stem from B-Type KRs and active DHs. The other three olefins present in the structure of palmerolide A, as noted above, likely have positional and stereochemical influence during the enzymatic shifts to the β,γ-positions (Δ^21^ and Δ^23^) or from the ECH domain (module 6).

Additional insights into the structural features of the resulting compound were obtained through defining the specificity of KS domains using phylogenetic analysis and the *trans*-AT PKS Polyketide Predictor (*trans*ATor) bioinformatic tool (Helfrich et al., 2019). KS domains catalyze the sequential two-carbon elongation steps through a Claisen-like condensation with a resulting β-keto feature (Khosla et al., 2007). Additional domains within a given module can modify the β-carbonyl or add functionality to the adjacent α- or γ-positions (Keatinge-Clay, 2012). Specificity of KSs, based on the types of modification located on the upstream acetate subunit were determined and found to be mostly consistent with our retrobiosynthetic predictions (**Supplemental Figure S4, Supplemental Table S1**). For example, the first KS, KS1 (module 1), is predicted to receive a subunit containing a β-branch. KS3 (module 4) and KS4 (module 5) are predicted to receive an upstream monomeric unit with α-methylation and an olefinic shift, consistent with the structure of palmerolide A and with the enzymatic transformations resulting from module 3 and module 4, respectively. Interestingly, the KS associated with the HCS cassette branches deeply compared to all others upon phylogenetic analysis (**Supplemental Figure S4**). *Trans*ATor also aided in confirming the stereochemical outcomes of the hydroxy groups and olefins, which occur through reduction of the β-carbonyls. The predictions for the D-configured hydroxy groups were consistent with not only the presence of the LDD motif, indicative of type B-type KR as outlined above, but also with stereochemical determination based on the clades of the KS domains of the receiving modules, KS5 (module 6) and KS11 (module 12). They are also consistent with the structure of palmerolide A. The KS predictions, however, did not aid in confirming reduction of the upstream olefins for KS8 (module 9) and KS12 (module 13).

#### Additional *trans*-acting domains and domains between genes responsible for biosynthesis

A glycosyl transferase (*palP*) and lactone oxidase (*palH*) that are often associated with glycosylation of polyketides are encoded in the palmerolide A BGC following the AT domains and preceding the HCS cassette (**Figure 3**). Though glycosylated palmerolides have not been observed, glycosylation as a means of self-resistance in *Streptomyces* has been described and is a possible explanation for these observed domains (Quirós et al., 1998; Wencewicz, 2019). Glycosyl transferases are found in other macrolide- and non-macrolide-producing organisms as a means to inactivate hydroxylated polyketides (Jenkins and Cundliffe, 1991; Gourmelen et al., 1998). Though prokaryotic V-ATPases tend to be more structurally simple than those of eukaryotes, there is homology in the active sites of prokaryotic and eukaryotic V-ATPases making the pro-drug hypothesis for self-resistance reasonable in palmerolide A biosynthesis (Yokoyama and Imamura, 2005). The D-arabinono-1,4-lactone oxidase (*palH*) is a FAD-dependent oxidoreductase that likely works in concert with the glycosyltransferase. An ATP-binding cassette (ABC) transporter is encoded between the core biosynthetic genes and the genes for the *trans-*acting enzymes (Murray et al., 2021). This transporter, which has homology to SryD and contains the key nucleotide-binding domain GGNGSGKST, may be responsible for the translocation of the macrolide out of the cell, since it is housed within the BGC, it is likely under the same regulatory control. Additionally, a Band7 protein is encoded. Band7 proteins belong to a ubiquitous family of slipin or stomatin-like integral membrane proteins that are found in all kingdoms, yet lack assignment of function (Boehm et al., 2009). A domain of unknown function (DUF) is annotated; however, this particular DUF, DUF179, has no superfamily relatives in protein homology searches. Together these genes encoding potential macrolide glycosylation and transport functions may play a role in the bioactivity and export of palmerolide A from the producing organism. Future integrated studies will be needed to decipher the functions of these genes *in situ*.

### C. Multiple copies of the *pal* Biosynthetic Gene Cluster Explain Structural Variants in the Palmerolide Family

Careful assembly of the Ca. *S. palmerolidicus* MAG revealed the *pal* BGC was present in multiple copies (**Figure 4** and **Supplemental Figure S5-S7**) (Murray et al., 2021), evidenced by their independent anchoring loci within the MAG and supported by a five-fold increase in depth of coverage relative to the rest of the genome. The structural complexity of the multicopy BGCs represents a biosynthetic system that is similar to that found in *Ca*. Didemnitutus mandela, another ascidian-associated verrucomicrobium in the family *Opitutaceae* (Lopera et al., 2017). The five distinct Type-1 PKS BGCs with significant regions of overlap are likely responsible for much of the structural diversity in the family of palmerolides (Diyabalanage et al., 2006; Noguez et al., 2011) (**Figure 4**). Palmerolide A, which is the predominant secondary metabolite isolated from *Synoicum adareanum* (Murray et al., 2020), is hypothesized to arise from the BGC designated as *pal* BGC 4 with additional compounds also arising from this cluster. The other clusters designated as *pal* BGC 1, *pal* BGC 2, *pal* BGC 3, and *pal* BGC 5 and their potential biosynthetic products of each are described below. It is hypothesized that there are three levels of diversity introduced to create the family of palmerolides: (1) differences in the site of action for the *trans*-acting domains (with additional *trans*-acting domains at play as well), (2) promiscuity of the initial selection of the starter subunit, and (3) differences in the core biosynthetic genes with additional PKS domains or stereochemical propensities within a module.

**Figure 4.**
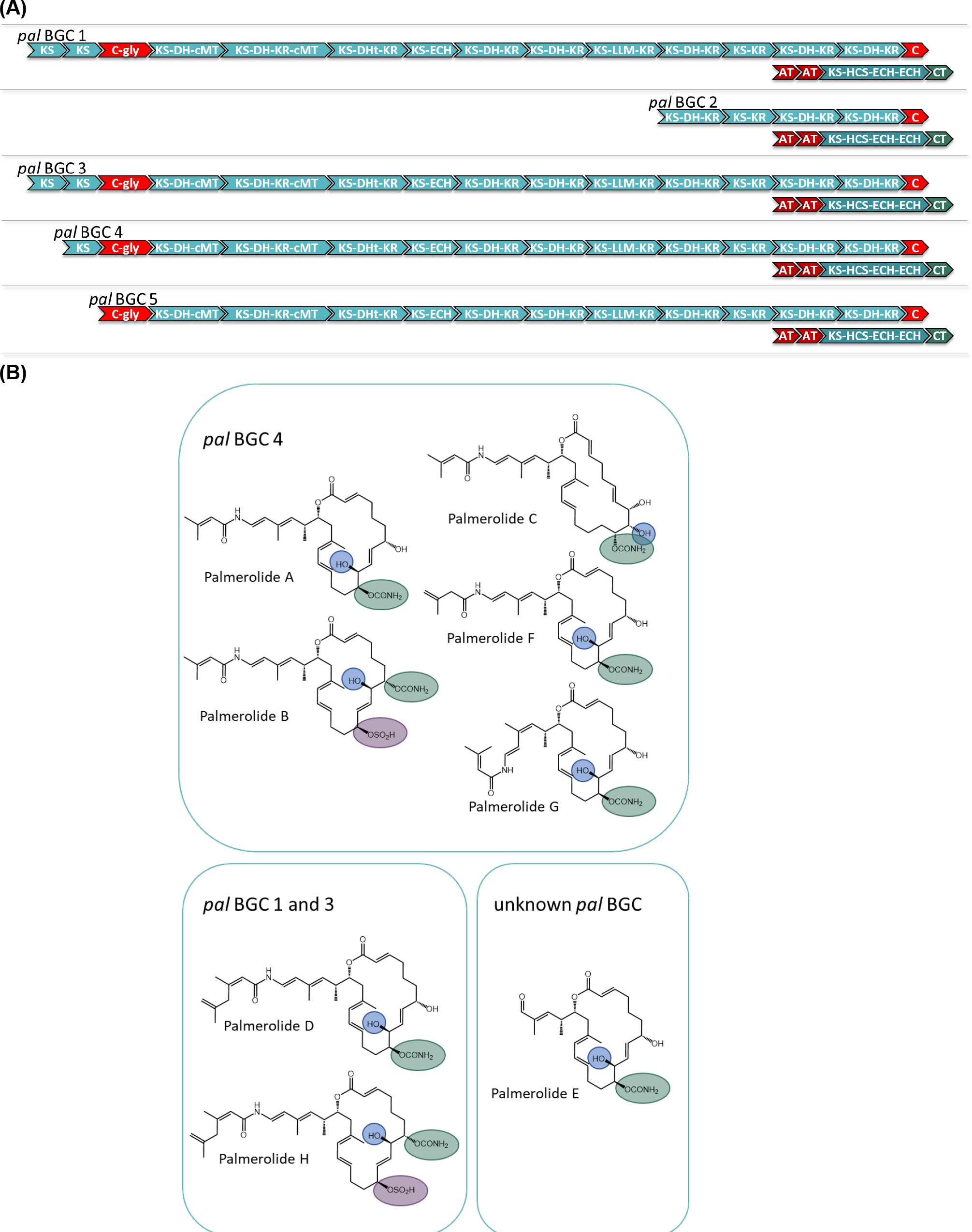
**(A)** Comparison of the modular structure of the 5 *pal* BGCs. **(B)** Family of palmerolides. Much of the structural diversity can be explained by differences due to starter unit promiscuity, sites of action for the *trans*-acting tailoring enzymes, and differences in the core modules of the multiple *pal* BGCs. It is proposed that *pal* BGC 4 is responsible for not only palmerolide A, but also palmerolide B, palmerolide C, palmerolide F, and palmerolide G. It is interesting to note that the modular structure of the domains responsible for biosynthesis are equivalent for *pal* BGC 1 and *pal* BGC 3. These two BGCs contain an additional KS domain as compared to *pal* BGC 4 and are likely responsible for the biosynthesis of palmerolide D and palmerolide H.

There are several palmerolides that likely arise from the same BGC encoding the megaenzyme responsible for palmerolide A (*pal* BGC 4). We hypothesize that the *trans*-acting domains have different sites of action than what is seen in palmerolide A biosynthesis. For example, the chemical scaffold of palmerolide B (**Figure 4**) is similar to palmerolide A, though the carbamate transfer occurs on the C-7 hydroxy group. Palmerolide B instead bears a sulfate group on the C-11 hydroxy group; proteins with homology to multiple types of sulfatases from the UniProtKB database (P51691, P15289, O69787, Q8ZQJ2) are found in the genome of *Ca*. S. palmerolidicus (Murray et al., 2021), but are not encoded within the BGCs. One of these *trans*-acting sulfatases likely modifies the molecule post-translationally. Other structural differences including the hydroxylation on C-8 instead of C-10 (as observed in palmerolide A) and the Δ^9^ olefin that differs from palmerolide A’s Δ^8^ olefin, are either due to a difference of the site of action of the LLM (module 9) or a *trans*-acting hydroxylase. Another member of the compound family, palmerolide C, has structural differences attributable to *trans*-acting enzymes as well. Again, a *trans*-acting hydroxylase or the LLM is proposed to be responsible for hydroxylation on C-8. A hydroxy group on C-9 occurs through reduction of the carbonyl. The carbamate installation occurs on C-11 after *trans*-acting hydroxylation or LLM hydroxylation. In addition, the Δ^8^ olefin in palmerolide A is not observed, but rather a Δ^6^ olefin.

Additional levels of structural variation are seen at the site of the starter unit, likely due to a level of enzymatic promiscuity of the second AT (PalF). This, combined with differences in the sites of action for the *trans*-acting domains, is likely responsible for the structural differences observed in palmerolide F (**Figure 4**). The terminal olefin on the tail of the macrolide, which perhaps is a product of promiscuity of the selection of the starter unit, the isomeric 3-methyl-3-butenoic acid, is consistent with the aforementioned lack of consensus for malonate selection by the AT. In addition, the KS that receives the starter unit is phylogenetically distinct from the other KS in the *pal* clusters (**Supplemental Figure S4**).

The retrobiosynthetic hypothesis for palmerolide G (**Figure 4**) has much similarity to what is present in *pal* BGC 4; however, the presence of a *cis*-olefin rather than a *trans*-olefin could arise from a difference in the enzymatic activity of module 4. This olefin subsequently undergoes an olefinic shift and, therefore, the stereochemistry is not solely reliant upon the action of the associated KR. Although this difference has not been identified in the BGCs in the samples sequenced, this could be present in other environmental samples that have been batched for processing and compound isolation. Currently, the biosynthetic mechanism is unknown.

The core biosynthetic genes of two palmerolide BGCs (*pal* BGC 1 and *pal* BGC 3) are identical to one another (**Supplemental Figure S5**) and possess an additional elongation module when compared to *pal* BGC 4. Palmerolide D (**Figure 4**) is structurally very similar to palmerolide A with the exception of elongation in the carboxylate tail of the macrolide by an isopropyl group. This could arise from one additional round of starter unit elongation via a KS and methylation. These two identical BGCs are consistent with the additional elongation module found in *pal* BGC1 and 3. The overall architecture and stereochemistry are otherwise maintained. Palmerolide H (**Figure 4**) also likely arises from these two BGCs although it includes the structural differences of both palmerolide B and palmerolide D in which it contains the extended carboxylate tail with a terminal olefin and incorporates hydroxylation on C-8 rather than C-10. Again, there is no genomic evidence that this hydroxylation in the α-position is due to incorporation of hydroxymalonate to explain this but is instead likely due to a *trans-*acting hydroxylase. The carbamate installation occurs on C-7, while sulfonation occurs on C-11 and α-hydroxy placement is on C-8.

The final two *pal* BGCs are shorter with a reduced number of biosynthetic modules found compared to *pal* BGC 4. The gene structure of *pal* BGC 5 (**Supplemental Figure S6**) shows preservation of many of the core biosynthetic genes; however, there are no pre-NRP PKS modules noted in the BGC. The HCS cassette, glycosyl transferase, and CT are all present downstream. The predicted product of this cluster does not correspond with a known palmerolide, though post-translational hydrolysis of the C-24 amide may result in a structure similar to palmerolide E (**Figure 4**), which maintains much of the structure of palmerolide A; however, it is missing the initial polyketide starter unit and the glycine subunit. The final *pal* BGC in *Ca*. S. palmerolidicus, *pal* BGC 2 (**Supplemental Figure S7**), includes only five elongating modules, which would result in a 10-carbon structure that has not been isolated. Interestingly, despite the shortened BGC, the HCS cassette, glycosyl transferase, and CT are all present downstream, and the sequence itself aligns perfectly with few SNPs to the other BGCs (Murray et al., 2021). There would only be a single hydroxy group serving as a substrate for the CT, glycosyl transferase, and sulfatase to act. The 2-carbon site of action for the β-branch introduced in the palmerolide A structures would not be present. The structure-based retrobiosynthesis of the eight known palmerolides (A-F) can be hypothesized to arise from differences in the core biosynthetic genes of these non-identical copies of the *pal* BGC, starter unit promiscuity, and differing sites of action in the *trans*-acting enzymes.

## Conclusion

The putative *pal* BGC has been identified and represents the first BGC elucidated from an Antarctic organism (Murray et al., 2021). As outlined in this retrobiosynthetic strategy, the *pal* BGC represents a *trans*-AT Type I PKS/NRPS hybrid system with compelling alignment to the predicted biosynthetic steps for palmerolide A. The *pal* BGC is proposed to begin with PKS modules resulting in the incorporation of an isovaleric acid derivative, 3-methylcrotonic acid, as a starter unit, followed by incorporation of a glycine residue with NRPS-type modules. Thereafter, eleven rounds of progressive polyketide elongation likely occur and leading to varying degrees of oxidation introduced with each module. There are several interesting non-canonical domains encoded within the BGC, such as an HCS, CT, LLM, and a truncated condensation termination domain. Additionally, a glycosylation domain may be responsible for reversible, pro-drug formation to produce self-resistance to the V-ATPase activity of palmerolide A. There are several additional domains, the function of which have yet to be determined.

A combination of modular alterations, starter unit differences, and activity of *trans*-acting enzymes contributes to Nature’s production of a suite of palmerolide analogues. There are a total of five distinct *pal* BGCs in the MAG of *Ca*. S. palmerolidicus, yielding the known eight palmerolides, with genetic differences that explain some of the structural variety seen within this family of compounds. These include differences in modules that comprise the core biosynthetic genes. Additionally, it is proposed that some of the architectural diversity of palmerolides arises from different sites of action of the *trans*-acting, or non-colinear, modules. Starter unit promiscuity is another potential source of the structural differences observed in the compounds. Analysis of the *pal* BGC not only provides insight into the architecture of this Type I PKS/NRPS hybrid BGC with unique features, but also lays the foundational groundwork for drug development studies of palmerolide A via heterologous expression.

### Data Deposition

Data referenced here was deposited to the NCBI under BioProject Accession Number PRJNA662631 (44). The GenBank Accession Number for the MAG of *Ca*. Synoicohabitans palmerolidicus is JAGGDC000000000 (44). The BGCs can be accessed in the MiBIG database with the associated Accession Numbers: BGC0002118 (for *pal* BGC 4) and BGC0002119 (for *pal* BGC 3).

## Author Contributions

This work was the result of a team effort in which the following contributions are recognized: conceptualization, A.E.M., P.S.G.C., and B.J.B.; methodology and experimentation, N.E.A., A.E.M., H.E.D., C.L., P.S.G.C., B.J.B; data analysis, N.E.A., A.E.M., H.E.D, C.L., K.W.D., A.E.K.D., P.S.G.C, B.J.B; data curation, N.E.A., A.E.M., H.E.D, C.L., P.S.G.C, B.J.B; original draft preparation, N.E.A., B.J.B., A.E.M., H.E.D.; review and editing, N.E.A., B.J.B., A.E.M., C.L., H.E.D., P.S.G.C., A.E.K.D; funding acquisition, A.E.M., P.S.G.C., and B.J.B. All authors have read and agreed to the published version of the manuscript.

## Acknowledgments

The authors acknowledge the assistance of field team members, including William Dent, Charles D. Amsler, James B. McClintock, Margaret O. Amsler, and Katherine Schoenrock. This work would not have been possible without the outstanding logistical support of the United States Antarctic Program. Lucas Bishop, Robert Read, and Mary L. Higham are also recognized for their contributions.

