## Supplemental Information for "Bioinformatic and mechanistic analysis of the palmerolide PKS-NRPS biosynthetic pathway from the microbiome of an Antarctic ascidian"

**Supplementary Figure 1.** Phylogenetic tree demonstrating the relationship between the third and fourth carrier proteins in the *pal* BGC with other carrier proteins ...............................pages 2-3

**Supplementary Figure 2.** Alignment of select carrier proteins in the *pal* BGC with select acyl-carrier proteins and peptidyl-carrier proteins ...........................................................................page 4

**Supplementary Figure 3. (A)** Alignment of the proposed palmerolide hydroxylase luciferase-like monooxygenase with sequences from the TIGR subfamily 04020. **(B)** Alignment of the proposed termination condensation domain of the *pal* BGC_4 with that of other related sequences ....page 5

**Supplementary Figure 4.** Phylogenetic comparison of the amino acid sequences of the KS Pfam domains from the *pal* BGCs compared to those from the BGCs for bryostatin, onnamide, pederin, psymberin, and sorangicin...........................................................................................pages 6-7

**Supplementary Figure 5.** The proposed BGC for *pal* BGC 1 .........................................pages 8-9

**Supplementary Figure 6.** The proposed BGC for *pal* BGC 5 .....................................pages 10-11

**Supplementary Figure 7.** The proposed BGC for *pal* BGC 2 .....................................pages 12-13

**Supplementary Table 1.** KS specificity predictions from *Trans*ATor.................................page 14

**Supplementary Table 2.** GenBank Accession Numbers and versions for the genomes including the BGCs used in Figure 3 and Figure 4. ..............................................................................page 15

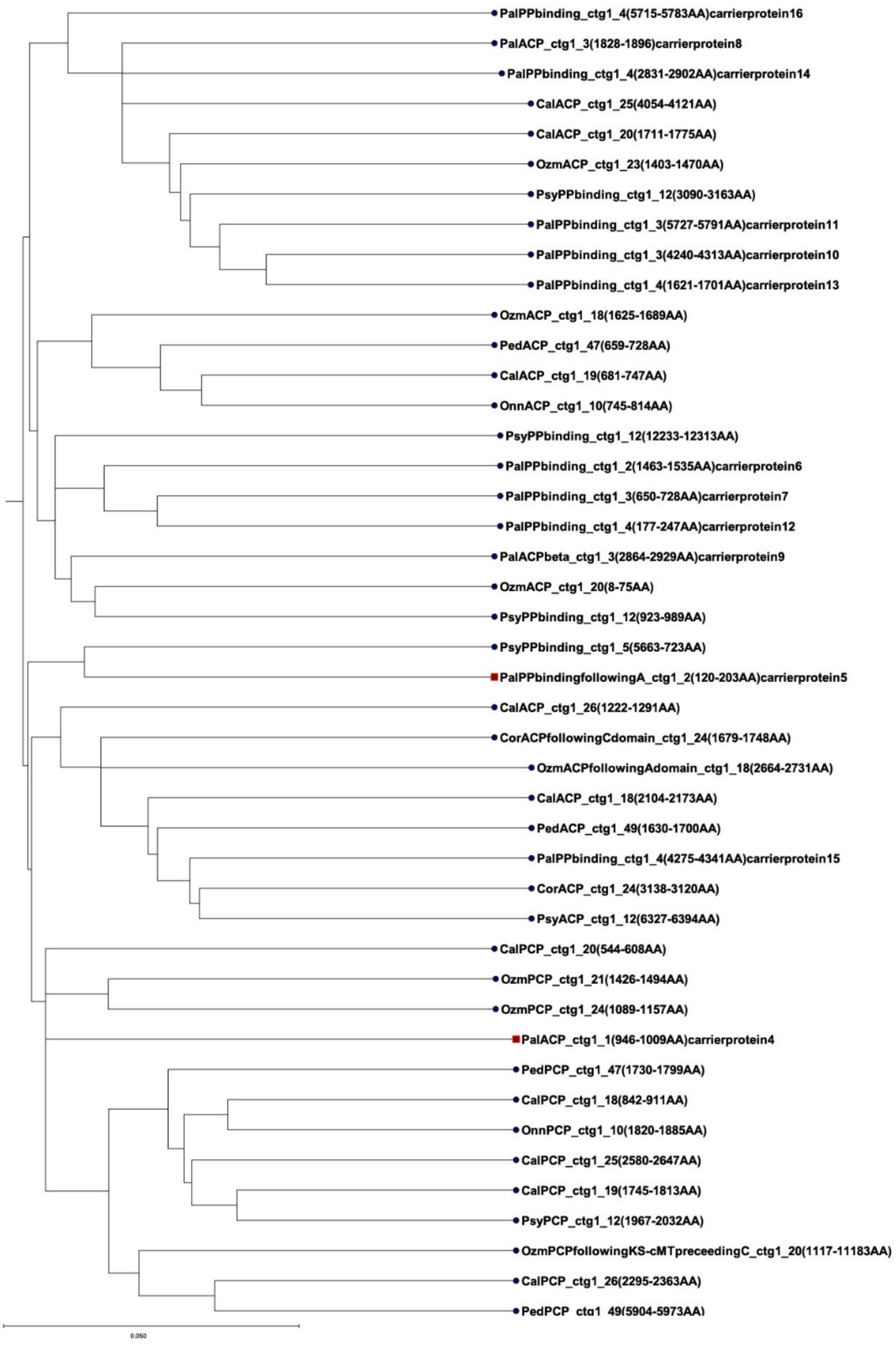
PalPPbinding_ctg1_4(5715-5783AA)carrierprotein16
PalACP_ctg1_3(1828-1896)carrierprotein8
PalPPbinding_ctg1_4(2831-2902AA)carrierprotein14
CalACP_ctg1_25(4054-4121AA)
CalACP_ctg1_20(1711-1775AA)
OzmACP_ctg1_23(1403-1470AA)
PsyPPbinding_ctg1_12(3090-3163AA)
PalPPbinding_ctg1_3(5727-5791AA)carrierprotein11
PalPPbinding_ctg1_3(4240-4313AA)carrierprotein10
PalPPbinding_ctg1_4(1621-1701AA)carrierprotein13
OzmACP_ctg1_18(1625-1689AA)
PedACP_ctg1_47(659-728AA)
CalACP_ctg1_19(681-747AA)
OnnACP_ctg1_10(745-814AA)
PsyPPbinding_ctg1_12(12233-12313AA)
PalPPbinding_ctg1_2(1463-1535AA)carrierprotein6
PalPPbinding_ctg1_3(650-728AA)carrierprotein7
PalPPbinding_ctg1_4(177-247AA)carrierprotein12
PalACPbeta_ctg1_3(2864-2929AA)carrierprotein9
OzmACP_ctg1_20(8-75AA)
PsyPPbinding_ctg1_12(923-989AA)
PsyPPbinding_ctg1_5(5663-723AA)
PalPPbindingfollowingA_ctg1_2(120-203AA)carrierprotein5
CalACP_ctg1_26(1222-1291AA)
CorACPfollowingCdomain_ctg1_24(1679-1748AA)
OzmACPfollowingAdomain_ctg1_18(2664-2731AA)
CalACP_ctg1_18(2104-2173AA)
PedACP_ctg1_49(1630-1700AA)
PalPPbinding_ctg1_4(4275-4341AA)carrierprotein15
CorACP_ctg1_24(3138-3120AA)
PsyACP_ctg1_12(6327-6394AA)
CalPCP_ctg1_20(544-608AA)
OzmPCP_ctg1_21(1426-1494AA)
OzmPCP_ctg1_24(1089-1157AA)
PalACP_ctg1_1(946-1009AA)carrierprotein4
PedPCP_ctg1_47(1730-1799AA)
CalPCP_ctg1_18(842-911AA)
OnnPCP_ctg1_10(1820-1885AA)
CalPCP_ctg1_25(2580-2647AA)
CalPCP_ctg1_19(1745-1813AA)
PsyPCP_ctg1_12(1967-2032AA)
OzmPCPfollowingKS-cMTpreceedingC_ctg1_20(1117-11183AA)
CalPCP_ctg1_26(2295-2363AA)
PedPCP_ctg1_49(5904-5973AA)
0.050

**Supplementary Figure 1.** Phylogenetic tree demonstrating the relationship between the third and fourth carrier proteins in the *pal* BGC with acyl-carrier proteins (ACPs) and peptidyl-carrier proteins (PCPs) from other hybrid PKS-NRPS systems. Though, an ACP would be expected at the end of module 1 (carrier protein 4) while a PCP would be expected at the end of module 2 (carrier protein 5), this is not what is observed bioinformatically. Carrier protein 4 was initially annotated as an ACP; however, is in the same clade as PCPs. Carrier protein 5 falls within the Pfam 00550.24 as a phosphopantetheine attachment site and is within the clade associated with ACPs.

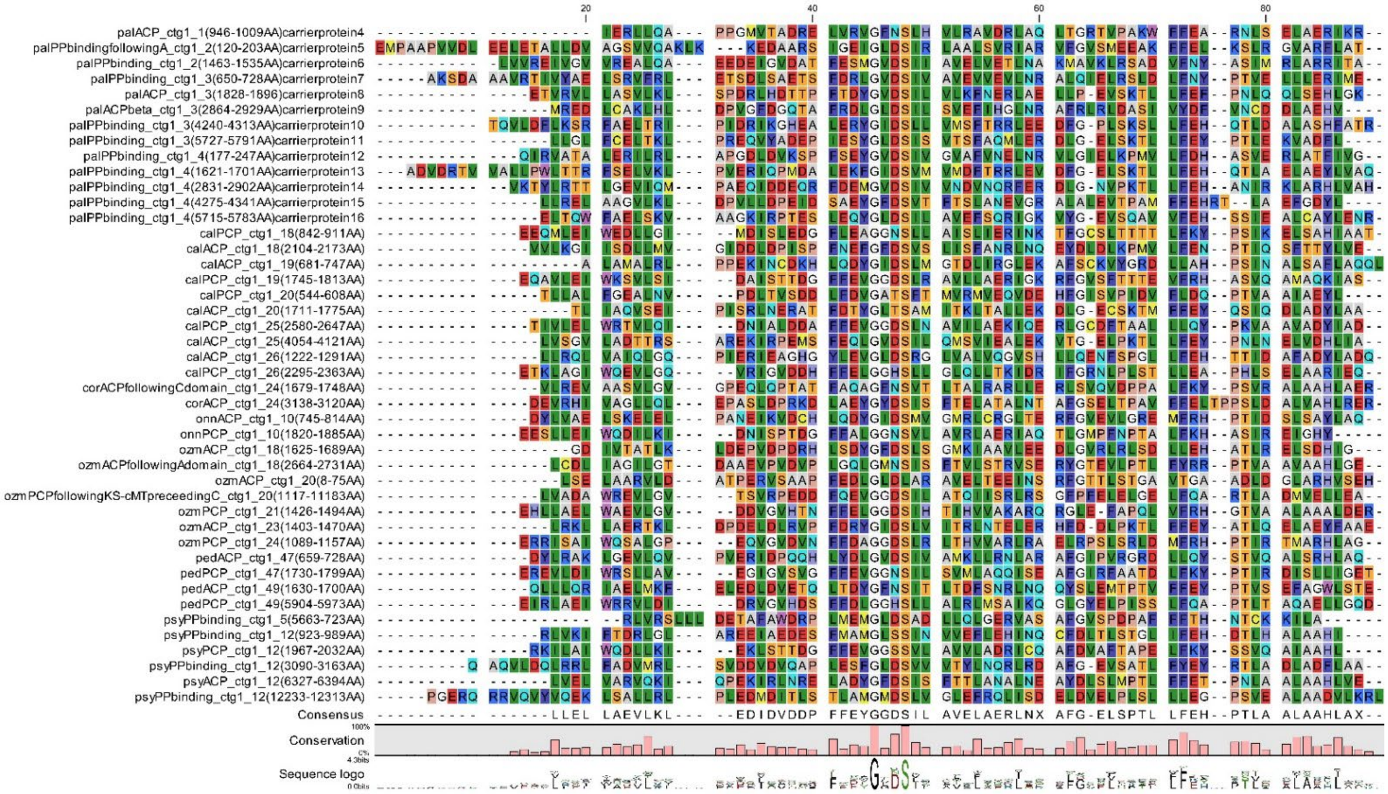

**Supplementary Figure 2.** Alignment of select carrier proteins in the *pal* BGC with acyl-carrier proteins (ACPs) and peptidyl-carrier proteins (PCPs) from other hybrid PKS-NRPS systems. The fifth carrier protein possess the (D/E)xGxDSL motif expected for a phosphopantetheine attachment site, though an isoleucine is present, rather than a leucine. This amino acid difference is not uncommon in other carrier proteins within PKS-NRPS BGCs.

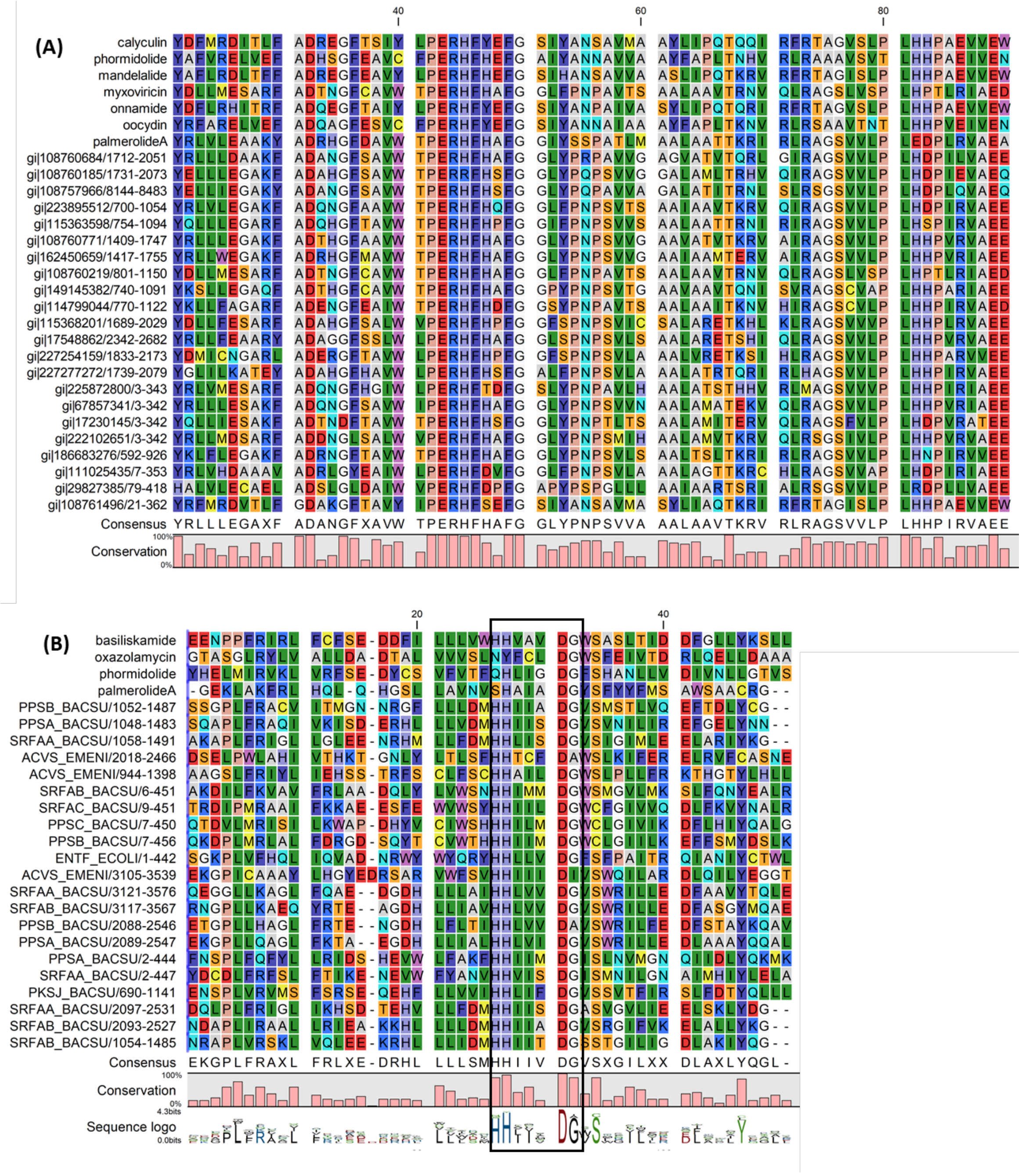

**Supplementary Figure 3. (A)** Alignment of the proposed palmerolide hydroxylase luciferase-like monooxygenase (LLM) with sequences from the TIGR subfamily 04020. **(B)** Alignment of the proposed termination condensation domain of the *pal* BGC_4 with that of basiliskamide and phormidolide as well as the HMM seed sequences for PF00668 (condensation domains); conservation of the HHXXXDG motif can be seen.

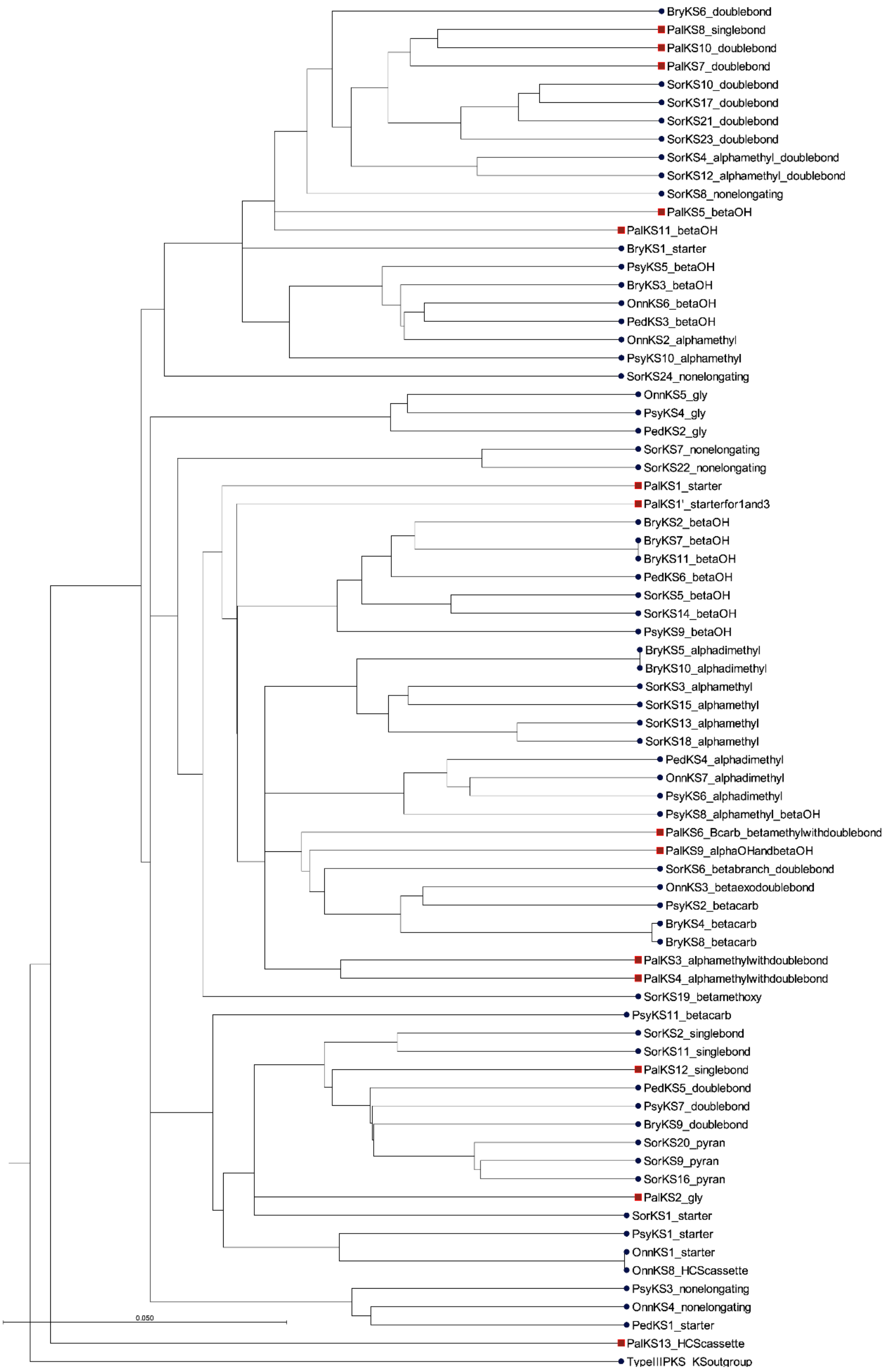
BryKS6_doublebond
PalKS8_singlebond
PalKS10_doublebond
PalKS7_doublebond
SorKS10_doublebond
SorKS17_doublebond
SorKS21_doublebond
SorKS23_doublebond
SorKS4_alphamethyl_doublebond
SorKS12_alphamethyl_doublebond
SorKS8_nonelongating
PalKS5_betaOH
PalKS11_betaOH
BryKS1_starter
PsyKS5_betaOH
BryKS3_betaOH
OnnKS6_betaOH
PedKS3_betaOH
OnnKS2_alphamethyl
PsyKS10_alphamethyl
SorKS24_nonelongating
OnnKS5_gly
PsyKS4_gly
PedKS2_gly
SorKS7_nonelongating
SorKS22_nonelongating
PalKS1_starter
PalKS1'_starterfor1and3
BryKS2_betaOH
BryKS7_betaOH
BryKS11_betaOH
PedKS6_betaOH
SorKS5_betaOH
SorKS14_betaOH
PsyKS9_betaOH
BryKS5_alphadimethyl
BryKS10_alphadimethyl
SorKS3_alphamethyl
SorKS15_alphamethyl
SorKS13_alphamethyl
SorKS18_alphamethyl
PedKS4_alphadimethyl
OnnKS7_alphadimethyl
PsyKS6_alphadimethyl
PsyKS8_alphamethyl_betaOH
PalKS6_Bcarb_betamethylwithdoublebond
PalKS9_alphaOHandbetaOH
SorKS6_betabranch_doublebond
OnnKS3_betaexodoublebond
PsyKS2_betacarb
BryKS4_betacarb
BryKS8_betacarb
PalKS3_alphamethylwithdoublebond
PalKS4_alphamethylwithdoublebond
SorKS19_betamethoxy
PsyKS11_betacarb
SorKS2_singlebond
SorKS11_singlebond
PalKS12_singlebond
PedKS5_doublebond
PsyKS7_doublebond
BryKS9_doublebond
SorKS20_pyran
SorKS9_pyran
SorKS16_pyran
PalKS2_gly
SorKS1_starter
PsyKS1_starter
OnnKS1_starter
OnnKS8_HCScassette
PsyKS3_nonelongating
OnnKS4_nonelongating
PedKS1_starter
0.050
PalKS13_HCScassette
TypeIIIPKS_KSoutgroup

**Supplementary Figure 4.** Phylogenetic comparison of the amino acid sequences of the KS Pfam domains from the *pal* BGCs compared to those from the BGCs for bryostatin, onnamide, pederin, psymberin, and sorangicin. The KS domains are numbered based on their order for proposed biosynthesis. Though many of the PalKSs are within their own subclades, they show homology with enzymes of similar substrate affinity. PalKS1 and PalKS1' (from *pal* BGCs 1 and 3) fall within the same clade, though do not clade with the KSs accepting the starter unit in the BGCs for sorangicin, onnamide, and psymberin, and instead demonstrate more homology with KSs receiving methylated subunits. A *trans*-acting ER has been hypothesized to fully reduce the subunits containing C4-5 and C12-13 (module 12 and module 8), but interestingly, the phylogeny did not prove informative regarding the differentiation between KS modules receiving upstream olefins versus fully reduced subunits (PalKS8 and PalKS12). Additionally, PalKS9 is in a subclade that is distinct from those receiving reduced subunits with β-hydroxy groups and instead has greater homology to the KSs responsible for accepting β-functionalized subunits such as an β-exo double bond, β-branch, and β-branch with endo-double bond (PalKS6). This supports the mechanism of the LLM acting as an α-hydroxylase while the elongating polyketide chain is online the biosynthetic megaenzyme and prior to the activity of the downstream module. Interestingly, PalKS13 which associated with the HCS cassette, is in a clade of its own. Note, the KS domain from the type III PKS BGC responsible for 3-(2'-hydroxy-3'-oxo-4'-methylpentyl)-indole biosynthesis from *Xenorhabdus bovienii SS-2004* was used for an outgroup.

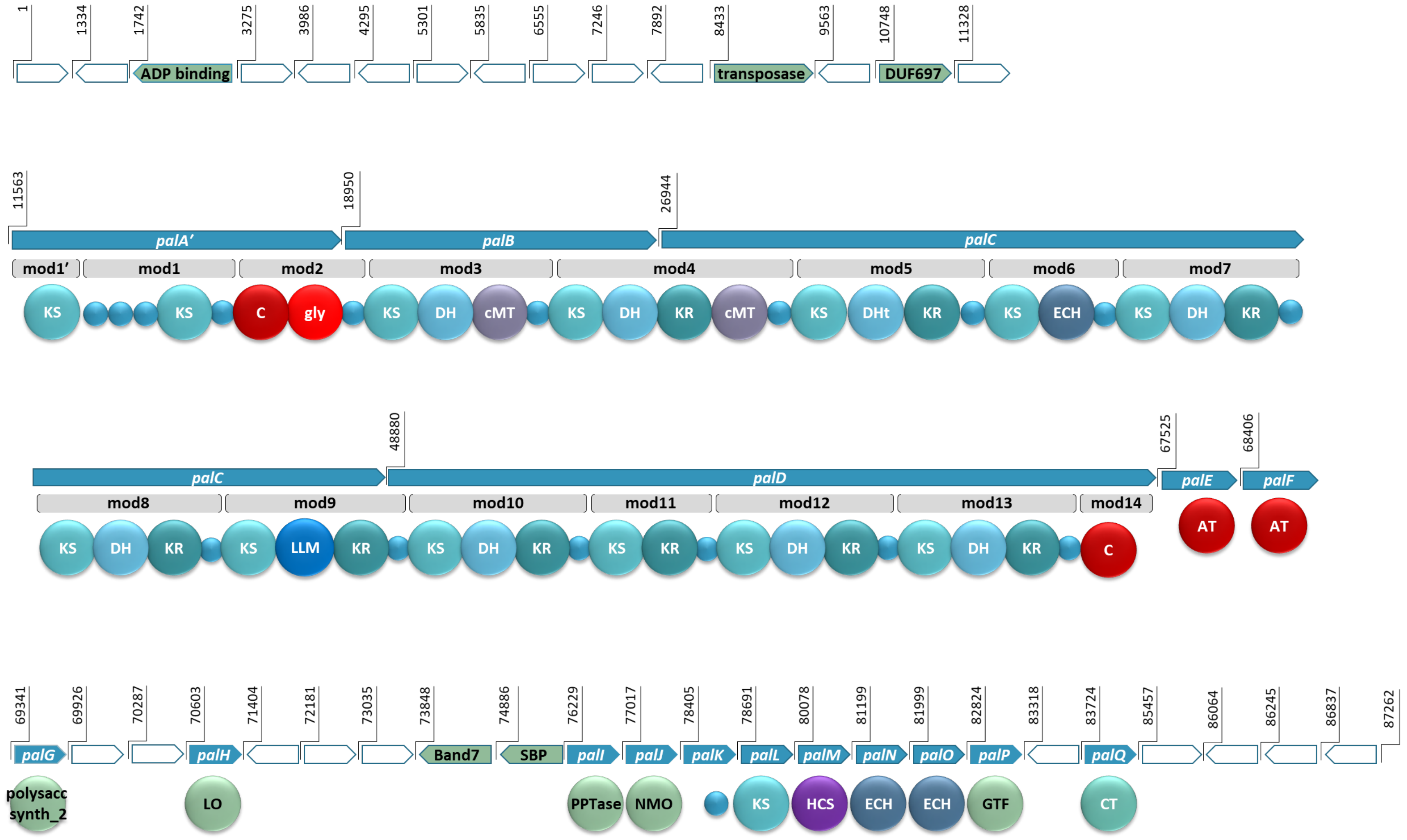

1
1334
1742
3275
3986
4295
5301
5835
6555
7246
7892
8433
9563
10748
11328
ADP binding
transposase
DUF697
11563
18950
26944
palA'
palB
palC
mod1'
mod1
mod2
mod3
mod4
mod5
mod6
mod7
KS
KS
C
gly
KS
DH
cMT
KS
DH
KR
cMT
KS
DHt
KR
KS
ECH
KS
DH
KR
48880
67525
68406
palC
palD
palE
palF
mod8
mod9
mod10
mod11
mod12
mod13
mod14
KS
DH
KR
KS
LLM
KR
KS
DH
KR
KS
KR
KS
DH
KR
KS
DH
KR
C
AT
AT
69341
69926
70287
70603
71404
72181
73035
73848
74886
76229
77017
78405
78691
80078
81199
81999
82824
83318
83724
85457
86064
86245
86837
87262
palG
palH
Band7
SBP
palI
palJ
palK
palL
palM
palN
palO
palP
palQ
polysacc synth_2
LO
PPTase
NMO
KS
HCS
ECH
ECH
GTF
CT

**Supplementary Figure 5.** The proposed BGC for *pal* BGC 1 which contains the same biosynthetic genes as *pal* BGC 3. The hybrid PKS-NRPS system has an additional elongation unit in *palA'* when compared to *palA,* seen at the beginning of the core biosynthetic genes. An ADP binding domain, a transposase, and a domain of unknown function (DUF697) are seen upstream of the core biosynthetic genes. KS: ketosynthase domain, C: condensation domain, gly: adenylation domain for glycine incorporation, DH: dehydratase domain, cMT: carbon methyl transferase domain, KR: ketoreductase domain, DHt: dehydratase variant, ECH: enoyl-CoA hydratase, LLM: luciferase-like monooxygenase, AT: acyltransferase; polysacc synt_2: polysaccharide biosynthesis protein, LO: lactone oxidase, ABC trans: ATP-binding cassette transporter, Band7: stomatin-like integral membrane protein, SBP: bacterial extracellular solute-binding protein, PPTase: phosphopantetheinyl transferase, NMO: nitronante monooxygenase, HCS: hydroxymethylglutaryl-CoA synthase, GTF: glycosyl transferase, CT: carbamoyl transferase, small blue circles represent acyl- or peptidyl-carrier proteins. Blue arrows indicate biosynthetic genes. Green arrows indicate genes that encode for non-biosynthetic proteins. White arrows reflect hypothetical genes. The BGC is displayed in reverse compliment.

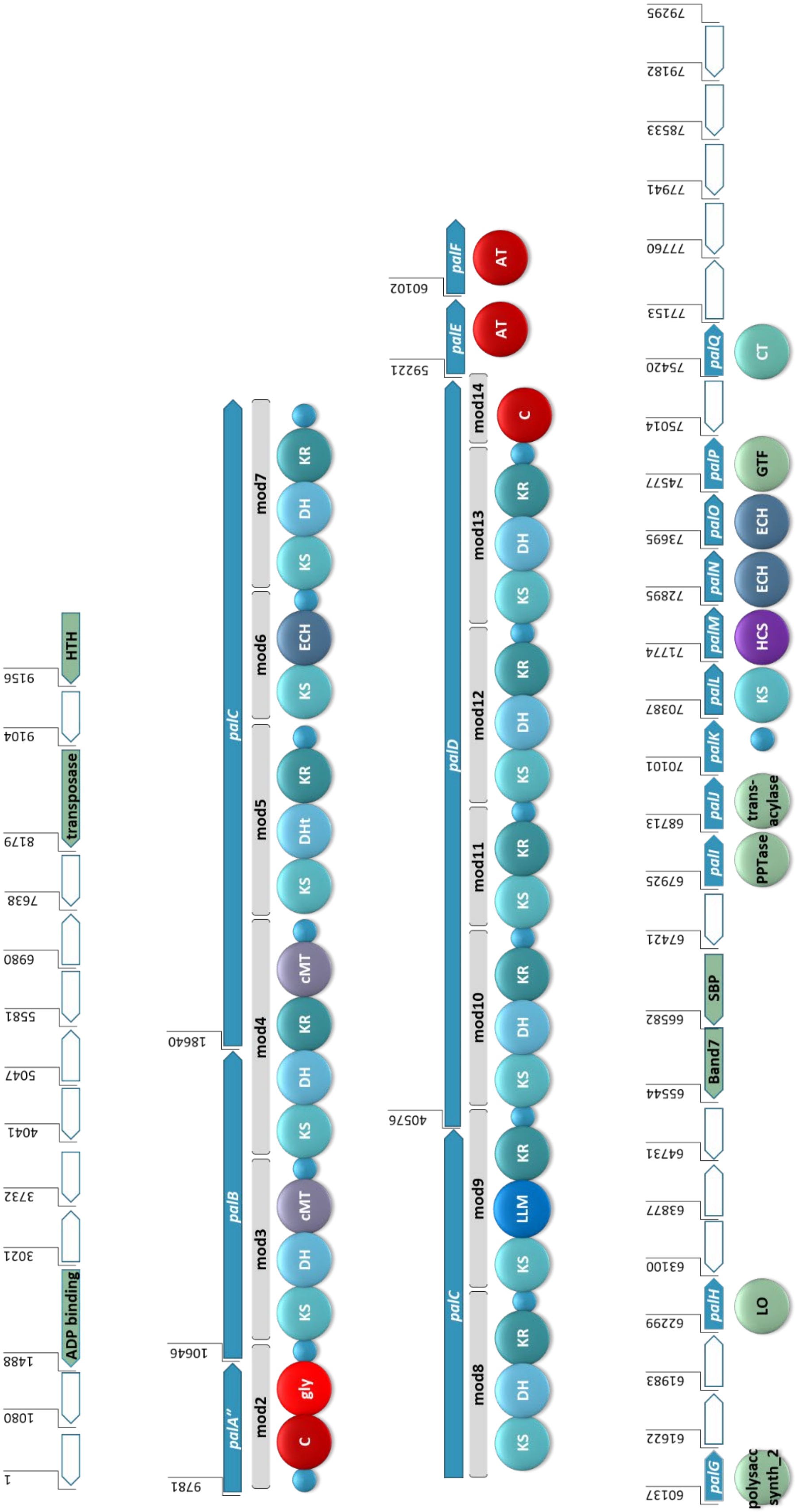

1
1080
1488
ADP binding
3021
3732
4041
5047
5581
6980
7638
8179
transposase
9104
9156
HTH
9781
10646
18640
palA''
palB
palC
mod2
mod3
mod4
mod5
mod6
mod7
C
gly
KS
DH
cMT
KR
DHt
ECH
40576
59221
60102
palC
palD
palE
palF
mod8
mod9
mod10
mod11
mod12
mod13
mod14
LLM
AT
60137
61622
61983
62299
63100
63877
64731
65544
66582
67421
67925
68713
70101
70387
71774
72895
73695
74577
75014
75420
77153
77760
77941
78533
79182
79295
palG
palH
palI
palJ
palK
palL
palM
palN
palO
palP
palQ
Band7
SBP
polysacc synth_2
LO
PPTase
trans-acylase
HCS
GTF
CT

**Supplementary Figure 6.** The proposed BGC for *pal* BGC 5, showing the hybrid PKS-NRPS system lacking the PKS portion of *palA* and designated here as *palA"* with the preservation of the enzymes responsible for incorporation of the glycine subunit. An ADP binding domain, a transposase, and a domain of unknown function (DUF697) are seen upstream of the core biosynthetic genes. KS: ketosynthase domain, C: condensation domain, gly: adenylation domain for glycine incorporation, DH: dehydratase domain, cMT: carbon methyl transferase domain, KR: ketoreductase domain, DHt: dehydratase variant; ECH: enoyl-CoA hydratase, LLM: luciferase-like monooxygenase, AT: acyl transferase; polysacc synt_2: polysaccharide biosynthesis protein, LO: lactone oxidase, ABC trans: ATP-binding cassette transporter, Band7: stomatin-like integral membrane, SBP: bacterial extracellular solute-binding protein, PPTase: phosphopantetheinyl transferase, NMO: nitronate monooxygenase, HCS: hydroxymethylglutaryl-CoA synthase, GTF: glycosyl transferase, CT: carbamoyl transferase, small blue circles represent acyl- or peptidyl-carrier proteins. Blue arrows indicate biosynthetic genes. Green arrows indicate genes that encode for non-biosynthetic proteins. White arrows reflect hypothetical genes. The BGC is displayed in reverse compliment.

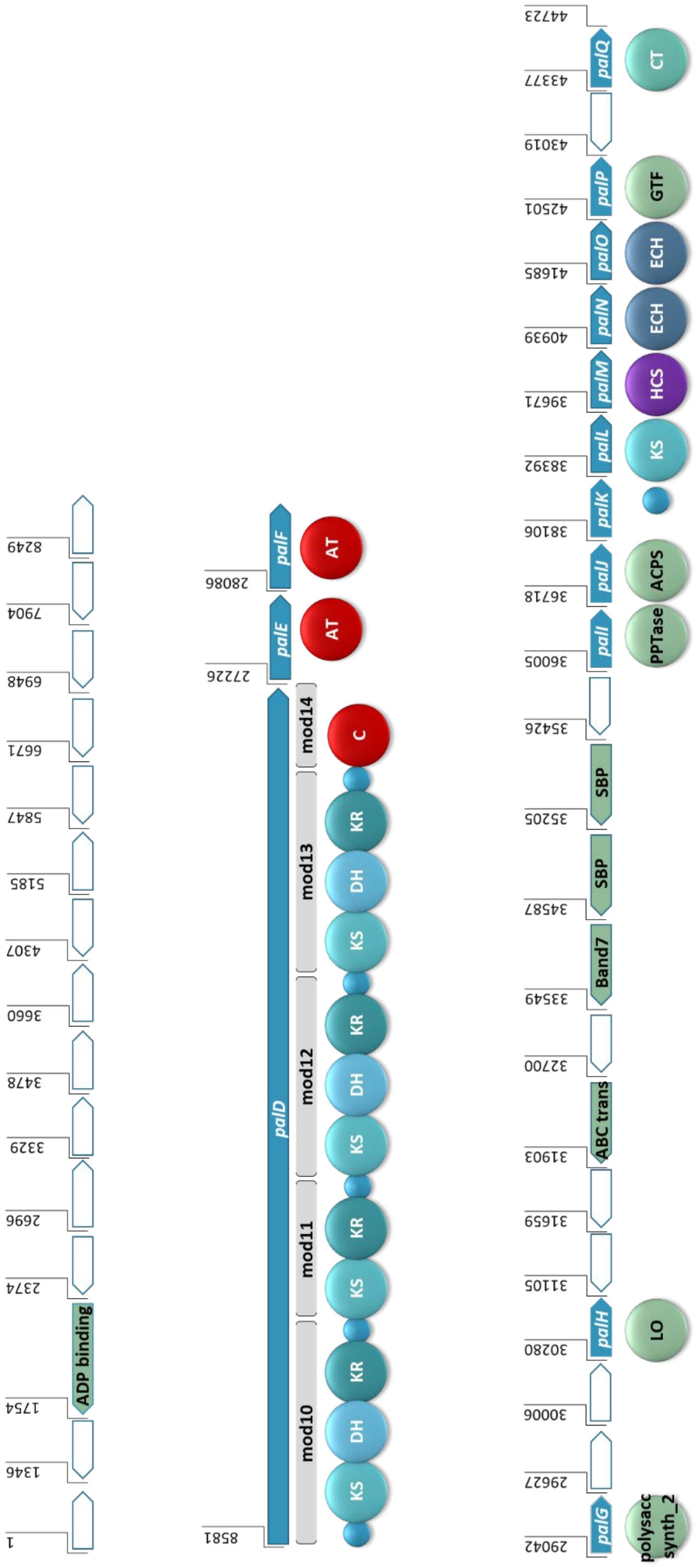

1
1346
1754
ADP binding
2374
2696
3329
3478
3660
4307
5185
5847
6671
6948
7904
8249
8581
palD
mod10
mod11
mod12
mod13
mod14
KS
DH
KR
KS
KR
KS
DH
KR
KS
DH
KR
C
27226
palE
AT
28086
palF
AT
29042
palG
polysacc synth_2
29627
30006
30280
palH
LO
31105
31659
31903
ABC trans
32700
33549
Band7
34587
SBP
35205
SBP
35426
36005
palI
PPTase
36718
palJ
ACPS
38106
palK
38392
palL
KS
39671
palM
HCS
40939
palN
ECH
41685
palO
ECH
42501
palP
GTF
43019
43377
palQ
CT
44723

**Supplementary Figure 7.** The proposed BGC for *pal* BGC 2, showing the shortened BGC with no NRPS portion, seen at the beginning of the core biosynthetic genes. An ADP binding domain is upstream of the core biosynthetic genes. The downstream tailoring enzymes are present, despite this cluster having the biosynthetic capability to only form an 8-carbon polyketide chain. KS: ketosynthase domain, C: condensation domain, DH: dehydratase domain, KR: ketoreductase domain, ECH: enoyl-CoA hydratase, AT: acyl transferase; polysacc synt_2: polysaccharide biosynthesis protein, LO: lactone oxidase, ABC trans: ATP-binding cassette transporter, SBP: bacterial extracellular solute-binding protein, PPTase: phosphopantetheinyl transferase, ACPS: acyl carrier protein synthase; HCS: hydroxymethylglutaryl-CoA synthase, GTF: glycosyl transferase, CT: carbamoyl transferase, small blue circles represent acyl- or peptidyl-carrier proteins. Blue arrows indicate biosynthetic genes. Green arrows indicate genes that encode for non-biosynthetic proteins. White arrows reflect hypothetical genes. The BGC is displayed in reverse compliment.

**Supplementary Table 1.** KS specificity predictions from *Trans*ATor. KS number designation is based on position in the pal BGC.

| | **Predicted specificity** | ***Trans*ATor Clade** | ***Trans*ATor Clade specificity** | **e-value** | **score** |
|---|---|---|---|---|---|
| **KS_1** | β-branch with Double Bond | 104 | β-O-Me or β-Me double bond | 1.10E-140 | 462 |
| **KS_2** | Gly_Single Bond | 96 | various specificities | 5.00E-171 | 562 |
| **KS_3** | Double Bond with -Me | 2 | α-Me shifted double bond or OH | 3.20E-189 | 622 |
| **KS_4** | Double Bond with -Me | 2 | α-Me shifted double bond or OH | 6.50E-191 | 627 |
| **KS_5** | Secondary D-OH | 53 | β-D-OH | 1.50E-192 | 633 |
| **KS_6** | B-carb, β-meth with Double Bond | 2 | α-Me shifted double bond or OH | 9.70E-192 | 630 |
| **KS_7** | Double Bond | 101 | double bonds | 4.20E-212 | 697 |
| **KS_8** | Single Bond | 101 | double bonds | 4.40E-214 | 704 |
| **KS_9** | Secondary -OH | 104 | β-O-Me or β-Me double bond | 9.50E-192 | 630 |
| **KS_10** | Double Bond | 127 | double bonds (E-config) | 1.40E-209 | 689 |
| **KS_11** | Secondary D-OH | 53 | β-D-OH | 8.90E-198 | 650 |
| **KS_12** | Single Bond | 96 | various specificities | 1.20E-197 | 650 |
| **KS_13** | HCS Cassette | 79 | starters or β-OH | 6.30E-36 | 117 |

**Supplementary Table 2.** GenBank Accession Numbers and versions for the genomes including the BGCs used in Figure 3 and Figure 4.

| Compound | BGC name | GenBank Accession | Version |
|---|---|---|---|
| **basiliskamide** | *bas* | NZ_AXBT01000013 | NZ_AXBT01000013.1 |
| **bryostatin 1** | *bry* | EF032014 | EF032014.1 |
| **calyculin** | *cal* | AB933566 | AB933566.1 |
| **corallopyronin** | *cor* | HM071004 | HM071004.1 |
| **mandelalide** | *mnd* | NJAL01000001 | NJAL01000001.1 |
| **onnamide** | *onn* | AY688304 | AY688304.2 |
| **oocydin** | *ooc* | JX315604 | JX315604.1 |
| **oxazolamycin** | *ozm* | EF552687 | EF552687.1 |
| **pederin** | *ped* | AH013687 | AH013687.2 |
| **phormidolide** | *phm* | KT727016 | KT727016.1 |
| **psymberin** | *psy* | FJ823461 | FJ823461.1 |
| **sorangicin** | *sor* | HM584908 | HM584908.1 |
| **myxoviricin** | *ta* | NC_008095 | NC_008095.1 |
